# Interoception of breathing and its relationship with anxiety

**DOI:** 10.1101/2021.03.24.436881

**Authors:** Olivia K. Harrison, Laura Köchli, Stephanie Marino, Roger Luechinger, Franciszek Hennel, Katja Brand, Alexander J. Hess, Stefan Frässle, Sandra Iglesias, Fabien Vinckier, Frederike Petzschner, Samuel J. Harrison, Klaas E. Stephan

## Abstract

Interoception, the perception of internal bodily states, is thought to be inextricably linked to affective qualities such as anxiety. While interoception spans sensory to metacognitive processing, it is not clear whether anxiety is differentially related to these processing levels. Here we investigated this question in the domain of breathing, using computational modelling and high-field (7 Tesla) fMRI to assess brain activity relating to dynamic changes in inspiratory resistance of varying predictability. Notably, the anterior insula was associated with both breathing-related prediction certainty and prediction errors, suggesting an important role in representing and updating models of the body. Individuals with low vs. moderate anxiety traits showed differential anterior insula activity for prediction certainty. Multimodal analyses of data from fMRI, computational assessments of breathing-related metacognition, and questionnaires demonstrated that anxiety-interoception links span all levels from perceptual sensitivity to metacognition, with strong effects seen at higher levels of interoceptive processes.

## Introduction

We perceive the world through our body. While questions regarding how we sense and interpret our external environment (exteroception) have been highly prominent across centuries of research, the importance and cognitive mechanisms of monitoring our internal environment have only more recently gained traction within the neuroscience community (Barrett and Simmons, 2015; Craig, 2002; Seth, 2013; Tsakiris and Critchley, 2016). ‘Interoception’, the perception of our body and inner physiological condition (Seth, 2013), constitutes a fundamental component of cerebral processes for maintaining bodily homeostasis (Berntson and Khalsa, 2021; Chen et al., 2021; Pezzulo et al., 2015; Quigley et al., 2021; Stephan et al., 2016). However, it has also been suggested to play a wider role within systems governing emotion, social cognition and decision making (Adolfi et al., 2017; Tsakiris and Critchley, 2016). An impaired ability to monitor bodily signals has also been hypothesised to exist across a host of psychiatric illnesses (Bonaz et al., 2021; Khalsa et al., 2017), and in particular for anxiety (Paulus, 2013; Paulus and Stein, 2010). As sympathetic arousal is a reflexive response to a perceived threat, many symptoms associated with anxiety manifest themselves in the body (such as a racing heart or shortness of breath). Conversely, perceiving bodily states compatible with sympathetic arousal in the absence of external triggers can itself induce anxiety (Paulus, 2013). Miscommunications between the brain and body are thus thought to represent a key component of anxiety, where bodily sensations may be under-, over- or mis-interpreted (Paulus and Stein, 2010), which can initiate and perpetuate symptoms of anxiety.

Studying interoception is not without significant challenges, as bodily signals are both noisy and difficult to safely manipulate (Khalsa et al., 2017). Controlled manipulations of respiratory processes represent a promising way to address these challenges: suitable experimental setups allow for dynamic yet safe changes in visceral aspects of respiration as one interoceptive modality (Berner et al., 2017; DeVille et al., 2018; Faull and Pattinson, 2017; Faull et al., 2016, 2018; Hayen et al., 2017; Paulus et al., 2012; Rieger et al., 2020). Furthermore, given the vitally important role of breathing for survival, respiratory changes are highly salient. Indeed, laboured, unsatisfied, unexpected or uncontrolled breathing can itself be perceived as a dangerous and debilitating interoceptive threat (Hayen et al., 2013; Herigstad et al., 2011; Schwartzstein et al., 1990). Beyond respiratory diseases (Carrieri-Kohlman et al., 2009; Hayen et al., 2013; Herigstad et al., 2011; Janssens et al., 2011; Marlow et al., 2019; Parshall et al., 2012), aversive breathing symptoms have been noted to be particularly prevalent in individuals suffering from psychiatric conditions such as anxiety and panic disorder (Giardino et al., 2010; Mallorqui-Bague et al., 2016; McNally and Eke, 1996; Paulus, 2013; Smoller et al., 1996; Woods et al., 1986).

Work towards conceptualising interoceptive dimensions has provided us with a framework to integrate the growing body of interoception research. Instead of treating interoception as a single entity, studies now consider both different sensory channels (e.g., organ-specific and humoral signals) and cognitive layers of interoceptive processing (Critchley and Garfinkel, 2017). These layers encompass multiple levels, ranging from metrics of afferent signal strength at ‘lower’ levels (using techniques such as heartbeat evoked potentials) (Allen et al., 2016; Petzschner et al., 2019) and psychophysical properties (such as measuring perceptual sensitivity (Domschke et al., 2010; Kleckner et al., 2015; Petzschner et al., 2017)) to psychological and cognitive components at ‘higher’ levels (Critchley and Garfinkel, 2017). Notable domains within these higher levels include attention toward bodily signals (Berner et al., 2017; Murphy et al., 2019; Wang et al., 2019), static and dynamic beliefs and models of body state (Critchley and Garfinkel, 2017; Seth, 2013; Tsakiris and Critchley, 2016), and insight into both our interoceptive abilities (Garfinkel et al., 2015, 2016b, 2016a; Harrison et al., 2020a) and the accuracy of our interoceptive beliefs (‘metacognition’) (Petzschner et al., 2017; Stephan et al., 2016). Importantly, research into dynamic models of body state has also connected the interoceptive literature to that of learning, where influential (Bayesian) theories of inference about the external world, e.g. predictive coding (Behrens et al., 2007; Feldman and Friston, 2010; Friston, 2005; O’Reilly and Jbabdi, 2012), have been extended to interoception and used to propose how brains may build models of the changing internal environment (Barrett and Simmons, 2015; Gu et al., 2013; Seth, 2013; Seth et al., 2012; Stephan et al., 2016). While theoretical models have been proposed, realistic synthetic data have been produced (Allen et al., 2019; Tschantz et al., 2021), and initial learning models have now been fit to empirical cardiac data (Smith et al., 2020), concurrent measures of dynamic brain processes during interoceptive learning have not yet been demonstrated.

Here, we build on these conceptual advances and assess the relationship between anxiety and breathing-related interoception across the multiple hierarchical levels of processing. Importantly, whilst many theoretical proposals have been put forward as to how anxiety may interrupt the brain’s processing of dynamic (trial-by-trial) interoceptive predictions and/or prediction errors (Allen, 2020; Barrett and Simmons, 2015; Brewer et al., 2021; Paulus, 2013; Paulus and Stein, 2006, 2010; Paulus et al., 2019), these are as yet untested. Therefore, within a rigorous assessment profile we included neuroimaging of a novel breathing-related interoceptive learning paradigm, providing the first empirical insight into the brain activity associated with interoceptive predictions and prediction errors. Furthermore, it is not yet known how alterations in interoceptive learning may relate to previously identified relationships between anxiety and lower-level breathing sensitivity (Garfinkel et al., 2016a; Tiller et al., 1987), higher-level beliefs (Ewing et al., 2017; Garfinkel et al., 2016b; Mehling, 2016; Paulus and Stein, 2010) or metacognition (Harrison et al., 2020b). Therefore, we aimed to both assess how anxiety is related to dynamic interoceptive learning, and additionally provide a unifying perspective on anxiety and breathing-related interoception across the hierarchical levels of interoceptive processing. We adopted a multimodal experimental approach to investigate multiple levels of breathing-related interoceptive processing, including low-level perceptual sensitivity and related higher-level metacognition via the Filter Detection Task (FDT) (Harrison et al., 2020a), subjective interoceptive beliefs via questionnaires, and trial-by-trial interoceptive learning and related brain activity in a novel Breathing Learning Task (BLT). Both the FDT and trial-by-trial behavioural and functional magnetic resonance imaging (fMRI) data from the BLT were analysed with separate computational models. All tasks were performed by two matched groups of low and moderate anxiety individuals, allowing us to evaluate the relationship between anxiety and each level of breathing-related interoceptive processing across the hierarchy, from sensitivity to metacognition.

## Methods

### Participants

Thirty individuals (pre-screened online for MRI compatibility, right handedness, non-smoking status, and no history of major somatic or psychological conditions) were recruited into each of two groups, either with very low anxiety (score of 20-25 on the Spielberger State-Trait Anxiety Inventory (Spielberger et al., 1970); STAI-T), or moderate anxiety (score>=35 STAI-T). The resulting mean (±std) trait anxiety score for the low anxiety group was 23.2±1.8 and for the moderate anxiety group 38.6±4.6. Groups were matched for age and sex (15 females in each group), with mean (±std) ages of 25.4±3.9 and 24.2±5.0 years for low and moderate anxiety groups, respectively. Study numbers were based on a power calculation for a two-sided two-sample t-test with an α-level of 5% (moderate effect size d=0.5), where a power of 90% is achieved with 30 participants in each group. Behavioural data (not used in any other analyses) from an additional 8 participants served to determine model priors, with four participants (two females and two males) from each of the low and moderate anxiety groups. In this way, prior values could be drawn from a comparable group of participants as the main study sample. All participants signed a written, informed consent, and the study was approved by the Cantonal Ethics Committee Zurich (Ethics approval BASEC-No. 2017-02330).

Behavioural data for an additional group of 15 individuals (12 female) were collected for model validation purposes. These participants were not pre-selected based on anxiety values (mean±std trait anxiety = 38.9±12.5), but were screened for non-smoking status and no history of major somatic or psychological conditions. Participants were aged (mean±std) 23.1±5.6 years. All participants signed a written, informed consent, and the study was approved by the New Zealand Health and Disability Ethics Committee (HDEC) (Ethics approval 20/CEN/168).

### Procedural overview

Each participant completed three tasks over two testing sessions: a behavioural session that included questionnaires and a task probing interoceptive sensitivity and metacognition (the Filter Detection Task, or FDT), and a brain imaging session where the Breathing Learning Task (BLT) was paired with fMRI. Each of these tasks and analyses are described below, and all analyses were pre-specified in time-stamped analysis plans (https://gitlab.ethz.ch/tnu/analysis-plans/harrison_breathing_anxiety). The length of time between testing sessions (mean±std) was 4±3 days for all participants. Participants in the validation group completed the BLT in a behavioural session only.

### Questionnaires

The main questionnaire set employed was designed to firstly capture subjective affective measures, and secondly both general and breathing-specific subjective interoceptive beliefs. The assignment of participants to groups was based on the Spielberger Trait Anxiety Inventory (STAI-T) (Spielberger et al., 1970). Affective qualities that were additionally assessed included state anxiety (Spielberger State Anxiety Inventory; STAI-S (Spielberger et al., 1970)), symptoms that are part of anxiety disorder (Generalised Anxiety Disorder Questionnaire; GAD-7 (Spitzer et al., 2006)), anxiety sensitivity (anxiety regarding the symptoms of anxiety; Anxiety Sensitivity Index; ASI-3 (Taylor et al., 2007)), and symptoms of depression (Centre for Epidemiologic Studies Depression Scale; CES-D (Radloff, 1977)). To obtain self-reports of body awareness we used the Body Perception Questionnaire (BPQ) (Porges, 1995), while the Multidimensional Assessment of Interoceptive Awareness Questionnaire (MAIA) (Mehling et al., 2012) was used to measure positive and ‘mindful’ attention towards body symptoms. We also measured breathing-related catastrophising using the Pain Catastrophising Scale (PCS-B) (Sullivan et al., 1995), and breathing-related vigilance using the Pain Vigilance Awareness Questionnaire (PVQ-B) (McCracken, 1997) (in both questionnaires, the word ‘breathless’ or ‘breathlessness’ was substituted for ‘pain’). Finally, the following supplementary questionnaires were included to explore possible contributing factors (e.g. general positive and negative affect, resilience, self-efficacy and fatigue): Positive Affect Negative Affect Schedule (PANAS-T) (Watson et al., 1988), Connor-Davidson Resilience Scale (Connor and Davidson, 2003), General Self-Efficacy Scale (Schwarzer et al., 1997), Fatigue Severity Scale (FSS) (Krupp et al., 1989). The STAI-T and CES-D were completed online as part of the pre-screening process; all other questionnaires were completed in the behavioural session at the laboratory.

### Filter detection task

To systematically test properties of breathing perception and related metacognition, we utilised a perceptual threshold breathing task (the Filter Detection Task; FDT), described in detail elsewhere (Harrison et al., 2020a). The FDT was used to determine interoceptive perceptual sensitivity, decision bias, metacognitive bias (self-reported confidence) and metacognitive performance (congruency between performance and confidence scores) regarding detection of very small variations in an inspiratory load. In this task (outlined in Figure 2A), following three baseline breaths either an inspiratory load was created via the replacement of an empty filter with combinations of clinical breathing filters, or the empty filter was removed and restored onto the system (sham condition) for three further breaths. All filter changes were performed behind participants, out of their field of view. After each trial of six breaths, participants were asked to decide whether or not a load had been added, as well as reporting their confidence in their decision on a scale of 1-10 (1=not at all confident in decision, 10=extremely confident in decision). An adapted staircase algorithm was utilised to alter task difficulty until participants were between 60-85% accuracy (Harrison et al., 2020a), and 60 trials were completed at the corresponding level of filter load once the threshold level had been identified (using a ‘constant’ staircase procedure, as described by (Harrison et al., 2020a)). Respiratory threshold detection (Garfinkel et al., 2016b), metacognitive bias (Rouault et al., 2018) and interoceptive metacognitive performance (Harrison et al., 2020b) have previously been linked to anxiety symptomology.

### Breathing Learning Task

To measure behaviour and brain activity concerning the dynamic updating of interoceptive beliefs or expectations under uncertainty, a novel associative learning task was developed and employed during functional magnetic resonance imaging (fMRI). In this Breathing Learning Task (BLT), 80 trials were performed where on each trial two visual cues were paired with either an 80% or 20% chance of a subsequent inspiratory resistive load. Participants were explicitly told the probabilities that were being used, as well as that the cues were paired together – if one cue indicated an 80% chance of resistance then the other must indicate a 20% chance. Participants were also told that the cues could only ***swap*** their contingencies throughout the task, and could not act independently of each other. The number of trials was limited to 80 to ensure feasibility for participants. The number and structure of the cue contingency swaps (n=4) within the 80 trials was chosen as simulated data demonstrated both parameter recovery and model identifiability of the three candidate learning models (see Supplementary Figure 2 and ‘Quantification and Statistical Analysis’ details below).

The visual information for the task was presented through the VisualStim system (Resonance Technology, Northridge, CA, US). As outlined in Figure 3, participants were required to predict (via button press) whether they would experience a breathing resistance following the presentation of one of the cues. The visual cues were counter-balanced such that each was first matched with an 80% chance of resistance for half the participants, as well as counter-balancing of whether the answer ‘yes’ to the prediction question was presented on the left or right of the screen. Following this prediction and a short (2.5s) pause, a circle appeared on the screen to indicate the stimulus period (5s), where participants either experienced inspiratory resistance (70% of their maximal inspiratory resistance, measured in the laboratory, delivered via a PowerBreathe KH2; PowerBreathe International Ltd, Warwickshire, UK) or no resistance was applied. Rest periods of 7-9s were pseudo-randomised between trials.

For the inspiratory resistances we used a mechanical breathing system that allows for remote administration and monitoring of inspiratory resistive loads (for technical details on resistance administration see previous work (Rieger et al., 2020)). The cue presentations were balanced such that half of all trials delivered the inspiratory resistance. Following an initial stable period of 30 trials, the stimulus-association pairing was swapped four times during the remainder of the 80 trials (i.e., repeated reversals; Figure 3). The trial sequence was pseudorandom and fixed across subjects to ensure comparability of the induced learning process. Following every stimulus, participants were asked to rate ‘How difficult was it to breathe?”, on a visual analogue scale (VAS) from “Not at all difficult” to “Extremely difficult”. Immediately following the final trial of the task, participants were also asked to rate “How anxious were you about your breathing” on a VAS from “Not at all anxious” to “Extremely anxious”.

Two representations of trial-wise quantities were employed for subsequent analyses of data from this task. First, a computational model (see below) provided dynamic estimates of both predictions and prediction errors on each trial. Second, a standard categorical approach represented trial-by-trial whether the subjects’ prediction decisions indicated the anticipated presence or absence of an upcoming inspiratory resistance, as well as unsurprising (i.e. following correct predictions) and surprising (i.e. following incorrect predictions) respiratory stimuli. The latter results are presented in the Supplementary Material (Supplementary Figures 4 and 6).

### Statistical analysis overview

All analyses and hypotheses were pre-specified in an analysis plan (https://gitlab.ethz.ch/tnu/analysis-plans/harrison_breathing_anxiety). Each measure within each task modality was first compared between groups, and multiple comparison correction was applied within modalities. Pre-selection and comparisons across low or moderate anxiety scores allowed us to control for important factors such as age and sex, with equal numbers of men and women recruited into each group. Finally, a cross-modal analysis was performed on the key measures from each task. As the trait anxiety measure that was used to recruit participants into the separate groups was not included in this final analysis, we employed a correlation-based approach as the full spectrum of scores for each task were available.

### Questionnaire analysis

Group differences were tested individually for the 13 scores resulting from the 12 questionnaires, with all questionnaires included except the trait anxiety score that was used to screen participants and assign them to groups. The data that was used for group comparisons were first tested for normality (Anderson-Darling test, with p<0.05 rejecting the null hypothesis of normally distributed data), and group differences were determined using either two-tailed independent t-tests or Wilcoxon rank sum tests. For the questionnaires, Bonferroni correction for the 13 tests was applied, requiring p<0.004 for a corrected significant group difference. Results with p<0.05 not surviving correction are reported as exploratory for questionnaires as well as all other data. In a secondary exploratory step, group difference analyses were then conducted on the questionnaires’ subcomponent scores (22 scores); please see Supplementary Figure 1.

### FDT analysis

Breathing-related interoceptive sensitivity (i.e. perceptual threshold) was taken as the number of filters required to keep task performance between ∼60-85% accuracy. Both decision bias and metacognitive performance from the FDT were analysed using the hierarchical HMeta-d statistical model (Fleming, 2017), as previously described (Harrison et al., 2020a). This model firstly utilises signal detection theory (Stanislaw and Todorov, 1999) to provide single subject parameter estimates for task difficulty (d’; not analysed as performance is fixed between 60-85% by design) and decision bias (*c*, akin to over- or under-reporting the presence of resistance with values below and above zero, respectively), as well as using a hierarchical Bayesian formulation of metacognitive performance (Mratio, calculated by fitting metacognitive sensitivity meta-d’, then normalising by single subject values for d’). Finally, metacognitive bias was calculated as the average confidence scores across all analysed trials.

The four FDT parameters that were used for group comparison analyses were: sensitivity (filter number at perceptual threshold), decision bias (*c*), metacognitive bias (average confidence scores over threshold trials) and metacognitive performance (Mratio). These data were first tested for normality (Anderson-Darling test, with p<0.05 rejecting the null hypothesis of normally distributed data), and group differences were determined using either two-tailed independent t-tests or Wilcoxon rank sum tests. Importantly, as the single subject Mratio parameters were fit using a whole-group hierarchical model (with equal group numbers), standard statistical comparisons between groups for these parameters can also be performed. Hypothesised group differences were based on previous findings, where respiratory threshold detection level was hypothesised to be higher (Garfinkel et al., 2016a; Tiller et al., 1987), perceptual decisions biased towards ‘yes’ (or deciding the resistance was present; denoted by more negative values for *c*), metacognitive bias to be lower (Rouault et al., 2018) and interoceptive metacognitive performance to be lower (Harrison et al., 2020b) with greater anxiety. Bonferroni correction for the four tests was applied, requiring p<0.013 for a corrected significant group difference.

### BLT analysis

#### Model space

For the trial-by-trial analysis of behavioural data from the BLT, we considered three computational models that are routinely used for associative learning tasks. This included a Rescorla Wagner (RW) model (Equation 1) and 2 variants of the Hierarchical Gaussian Filter (HGF) with 2 or 3 levels (HGF2 and HGF3). While the RW model assumes a fixed learning rate, the HF allows for online adaption of learning rate as a function of volatility. All learning models were paired with a unit-square sigmoid response model (Equation 2) and were implemented using the Hierarchical Gaussian Filter Toolbox (Mathys et al., 2011, 2014) (version 5.3) from the open-source TAPAS software (Frässle et al.) (http://www.translationalneuromodeling.org/tapas/).

#### Prior selection and simulation analyses

Prior means and variances were determined using the distribution of maximum likelihood estimates fit across a holdout data set (consisting of 8 participants who were distinct from the participants of our study). Individual fits and estimated prior densities of the free parameters are given in Supplementary Table 2, as well as values for all remaining parameters of the three models. By adopting this procedure, prior densities were in a regime of the parameter space that is representative of the actual behavioural responses observed when participants performed the task. At the same time, the arbitrariness inherent to the specification of prior densities in non-hierarchical inference is reduced to a minimum.

To demonstrate face validity of the three models considered in our model space (each with one parameter free: *α* in RW, *ω*_2_ in HGF2, *κ*_2_ in HGF3), we assessed both parameter recovery and model identifiability for each (Wilson and Collins, 2019). Data for 60 synthetic subjects were generated for each of the candidate models by randomly sampling values from the prior densities that were placed over the parameters of the perceptual model. This synthetic data was generated for different noise levels (*ζ*_*sim*_ = 1,5,10). Subsequently, maximum a posteriori (MAP) parameter estimates were obtained using the Brayden-Fletcher-Goldfarb-Shanno algorithm, as implemented in the HGF toolbox, to fit the synthetic data sets. This allowed us to quantify parameter recovery and model identifiability across three different noise levels for each of the candidate models. Parameter recovery of the perceptual parameters was assessed using Pearson’s correlation coefficient (PCC) and by visual inspection of simulated and recovered parameter values (Supplementary Figure 2). Mean and standard deviation of estimated *ζ* values (from the response model) were computed for every noise level. Model identifiability was quantified by calculating the proportion of correctly identified models using approximate log model evidence (LME) scores and assessing whether the former was greater than the upper bound of the 90% confidence interval when assuming every model is equally likely a priori. For the resulting confusion matrices (Supplementary Figure 2), we additionally computed the mean proportion of correctly identified models (balanced accuracy scores).

The outlined procedure for assessing parameter recovery and model identifiability was repeated over 10 iterations with different seed values, to ensure robustness against any particular setting of the random number generator. The final results (PCCs for the perceptual parameters and *ζ*_*est*_ values) for every given level of noise were calculated as the average over all iterations, and are presented in Supplementary Table 3.

#### Model comparison and selection

Following MAP estimates of the 60 empirical datasets using each of the candidate models, the models were formally compared using random effects Bayesian model selection (BMS) as implemented in SPM12 (Friston et al., 2011; Rigoux et al., 2014; Stephan et al., 2009). BMS utilises the LME to determine the most likely amongst a set of competing hypotheses (i.e. models) that may have generated observed data, and is robust to outliers (Stephan et al., 2009). Our analysis plan had specified that a model would be chosen as the ‘winning’ model if it demonstrated a protected exceedance probability (PXP) greater than 90%. As explained in the Results section, none of our models reached this criterion (although simulations indicated that the proposed models could in principle be differentiated; see Supplementary Figure 2 for details). We therefore applied the simplest of the models considered (i.e. the RW model), as pre-specified in our analysis plan (https://gitlab.ethz.ch/tnu/analysis-plans/harrison_breathing_anxiety).

In our application of the RW model as a perceptual model, the update equation corresponded to a simple delta-learning rule with a single free parameter, the learning rate (Rescorla et al., 1972):

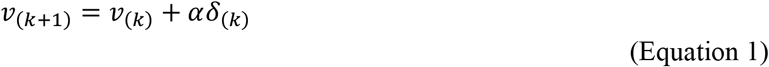

where *v*_(*k+1*)_ is the predicted probability for a specific outcome (encoded as 0 or 1) on trial (*k* + 1), *v*_(*k*)_ is the estimated outcome probability on the *k*^th^ trial, *α* ∈ [0, 1] is a constant learning rate parameter, and *δ*_(*k*)_ is the prediction error magnitude at trial *k*.

The above perceptual model was paired with a unit-square sigmoid response model (Mathys et al., 2014). This response model accounts for decision noise by mapping the predicted probability *v*_(*k*)_ that the next outcome will be 1 onto the probabilities *p*(*y*_(*k*)_ = 1) and *p*(*y*_(*k*)_ = 0) that the agent will choose response 1 or 0, respectively:

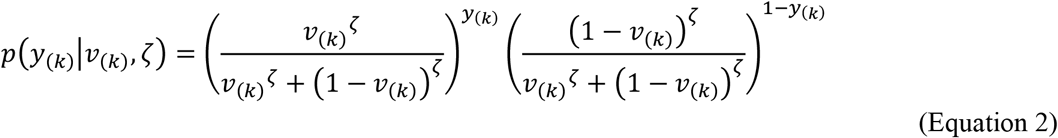

Here, *y*_(*k*)_ represents the expressed decision of a subject given the cue (contingency pairs) on trial *k*. The parameter *ζ* captures how deterministically *y* is associated with *v*. The higher *ζ*, the more likely the agent is to choose the option that is more in line with its current prediction. The decision model uses the perceptual model indirectly via its inversion (Mathys et al., 2014), given the trajectories of trial-wise cues and responses (see Figure 3).

In our paradigm, trial-wise outcomes are categorical (resistance vs. no resistance), which raises the question of how outcomes should be coded in the computational model. One way would be to model two trajectories, separately for resistance and no resistance outcomes, and indicate on any given trial whether the respective outcome has occurred (1) or not (0). However, due to the fixed coupling of contingencies in our paradigm (see above) – which the participants were explicitly instructed about – a computationally more efficient approach that requires only a single model is to code the outcome in relation to the cue. Here, we adopted this coding in “contingency space”, following the same procedure as in the supplementary material of Iglesias and colleagues (Iglesias et al., 2013). Specifically, due to the fixed coupling of contingencies in our paradigm (see above), we represented the occurrence of “no resistance” given one cue and the occurrence of “resistance” given the other cue as 1, and both other cue-outcome combinations as 0 (note that under the subsequent transformations described below, the resulting trajectories of predictions and prediction errors would remain identical if the opposite choice had been made).

#### Comparison of fitted model parameters

Group differences in model parameter estimates of learning rate (*α*) and inverse decision temperature (*ζ*), as well as perception measures of stimulus intensity (averaged across all trials), breathing-related anxiety (rated immediately following the task) and prediction response times were tested following tests for data normality. Bonferroni correction for five tests was applied, requiring p<0.01 for a corrected significant group difference. Results from additional exploratory models encompassing anxiety, depression and gender are reported in Supplementary Table 6.

#### Model validation

Following random effects Bayesian model selection (BMS (Rigoux et al., 2014; Stephan et al., 2009)), the chosen model was examined in each participant with regard to whether it demonstrated an adequate fit. To this end, model fit in each individual was compared to the likelihood of obtaining the data by a ‘null model’ (i.e. due to chance) (Daw, 2011) using the likelihood ratio test (*lratiotest* function) provided in MATLAB. The final behavioural and brain imaging analyses presented here were run without two subjects in which non-significant (p>0.05) differences to randomness were encountered. To demonstrate the extent to which the chosen model (the Rescorla Wagner) captured important aspects of participant performance, the proportion of incorrect responses across participants at each trial was compared to the mean prediction error trajectory (Supplementary Figure 3B).

To further validate the application of the chosen model to the BLT, we compared the fitted model trajectory to unseen behavioural data from an additional 15 participants. For qualitative assessment, the mean prediction error trajectory from the original dataset was compared to the proportion of incorrect responses across these held-out participants at each trial (Supplementary Figure 3C). For quantitative assessment, a logistic regression was conducted to assess whether the model prediction trajectory from the original data was able to significantly explain the prediction decisions made by the 15 participants in the validation sample.

#### Computationally informed regressors

The trajectories of predictions and prediction errors estimated by the RW model were used to construct regressors representing computational trial-by-trial quantities of interest for subsequent GLM analyses. In order to investigate the salient effects of inspiratory resistance as an interoceptive stimulus, we separated trials into “negative” (occurrence of resistance) and “positive” (no resistance) events and represented these events by separate regressors in the GLM (see Figure 4). To achieve this, we first transformed both the original prediction and prediction error values (estimated in contingency space) back into the stimulus space, according to the cue presented at each trial:

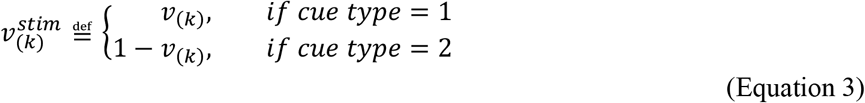

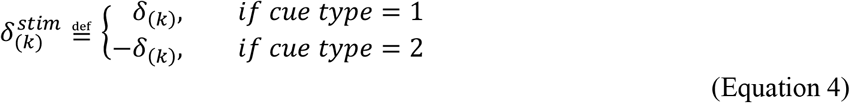

Here, 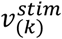 and 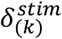 now represent the prediction and prediction error values in stimulus space, with 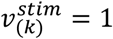 representing maximal predictions of no resistance and 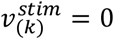 maximal predictions of resistance. Similarly, 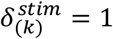 represents maximal prediction errors of no resistance and 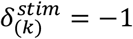 maximal prediction errors of resistance (see Supplementary Figure 4 for details).

Secondly, trial-wise prediction values were then transformed to represent the deviation from maximally uninformed predictions (i.e., guessing), by taking the distance from 0.5 (see Equations 5 and 6). In the RW model prediction values are probabilities bounded by 0 and 1, hence the distance from ‘guessing’ (at 0.5) reflects the ‘certainty’ by which the absence or presence of respiratory resistance was predicted. This simple transformation enabled us to take into account the role of (un)certainty of predictions – which plays a crucial role in interoception-oriented theories of anxiety (Paulus and Stein, 2010; Paulus et al., 2019) but, in contrast to Bayesian models, is not represented explicitly in the RW model. Specifically, separately for the two event types, we defined certainty of positive predictions (no resistance) and of negative predictions (resistance) as the absolute deviation from a prediction with maximum uncertainty (i.e., 0.5):

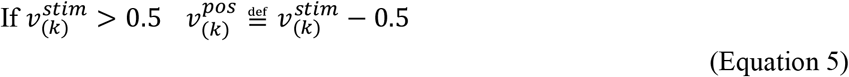

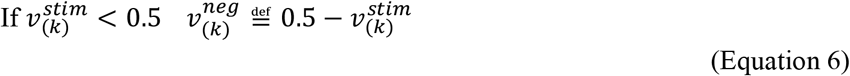

Here, both 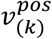 and 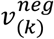 exist between 0 and 0.5, with values closer to zero indicating less certain predictions.

Like predictions, prediction errors were also divided between positive (no resistance) and negative events (resistance) values. This was again determined as the absolute deviation from the mid-point of the prediction errors (i.e., 0):

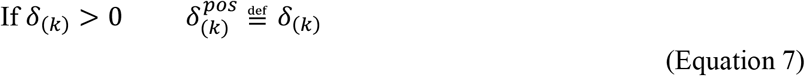

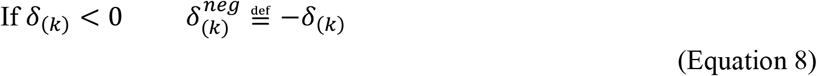

Here, both 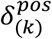 and 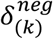 exist between 0 and 1, with values closer to zero indicating smaller prediction errors. Note that this derivation gives prediction error values identical to those that would have been obtained by modelling two separate trajectories for resistance and no resistance outcomes (see above and (Iglesias et al., 2013)).

#### Physiological data processing

Physiological data were recorded at a sampling rate of 1000 Hz, and included heart rate, chest distension, pressure of expired carbon dioxide (P_ET_CO_2_) and oxygen (P_ET_O_2_), and pressure at the mouth (for equipment details see (Rieger et al., 2020)). In addition to the task, small boluses of a CO_2_ gas mixture (20% CO_2_; 21% O_2_; balance N_2_) were administered during some rest periods, allowing for de-correlation of any changes in P_ET_CO_2_ from task-related neural activity, as previously described (Faull and Pattinson, 2017; Faull et al., 2015, 2016, 2018).

Physiological noise regressors were prepared for inclusion into single-subject general linear models (GLMs, described below). Linear interpolation between P_ET_CO_2_ peaks was used to form an additional CO_2_ noise regressor, which was convolved using a response function based on the haemodynamic response function (HRF) provided by SPM with delays of 10s and 20s for the overshoot and undershoot, respectively (Chang and Glover, 2009). Temporal and dispersion derivatives of this CO_2_ noise regressor were also included. An additional three cardiac- and four respiratory-related waveforms (plus one interaction term) were created using PhysIO (Kasper et al., 2017). Four respiratory volume per unit time (RVT) regressors (delays: -5, 0, 5, 10) were created using the Hilbert-transform estimator in PhysIO (Harrison et al., 2021), and convolution with a respiratory response function (Kasper et al., 2017).

#### Magnetic resonance imaging

MRI was performed using a 7 Tesla scanner (Philips Medical Systems: Achieva, Philips Healthcare, Amsterdam, The Netherlands) and a 32 channel Head Coil (Nova Medical, Wilmington, Massachusetts, United States of America). A T2*-weighted, gradient echo EPI was used for functional scanning, using a reduced field of view (FOV) with an axial-oblique volume centred over the insula and midbrain structures. The FOV comprised 32 slices (sequence parameters: TE 30ms; TR 2.3s; flip angle 75°; voxel size 1.5×1.5×1.5mm; slice gap 0.15mm; SENSE factor 3; ascending slice acquisition), with 860 volumes (scan duration 33 mins 9s). A matched whole-brain EPI scan (96 slices) was immediately acquired following the task scan for registration purposes. Additionally, a whole-brain T1-weighted structural scan with 200 slices was acquired (MPRAGE, sequence parameters: TE 4.6ms; TR 10ms; segment-TR 3000ms; TI 1000ms; flip angle 8°; voxel size 0.8×0.8×0.8mm; bandwidth; 153.1Hz/Px; sagittal slice orientation). Finally, a task-free (resting-state) functional scan (250 volumes) was obtained, with participants instructed to keep their eyes open and fixating a white fixation cross on a black screen.

#### MRI preprocessing

MRI data analysis was performed using a combination of FSL version 6.0.1 (the Oxford Centre for Functional Magnetic Resonance Imaging of the Brain Software Library, Oxford, UK) (Jenkinson et al., 2012) and SPM12 (Statistical Parametric Mapping software, London, UK) (Friston et al., 2011) as prespecified in our analysis plan (https://gitlab.ethz.ch/tnu/analysis-plans/harrison_breathing_anxiety). Image preprocessing was performed using FSL, including motion correction (MCFLIRT (Jenkinson and Smith, 2001)), removal of non-brain structures (BET (Smith, 2002)), and high-pass temporal filtering (Gaussian-weighted least-squares straight line fitting; 100s cut-off period)(Woolrich et al., 2001). Independent component analysis (ICA) was used to identify noise due to motion, scanner and cerebrospinal fluid artefacts (Griffanti et al., 2017), and the timeseries of these noise components were entered into single-subject GLMs (described below) as nuisance regressors. The functional scans were registered to the MNI152 (1×1×1mm) standard space using a three-step process: 1) Linear registration (FLIRT) with 6 degrees of freedom (DOF) to align the partial FOV scan to the whole-brain EPI image (Jenkinson et al., 2002); 2) Boundary-based registration (BBR; part of the FMRI Expert Analysis Tool, FEAT) with 12 DOF and a weighting mask of the midbrain and insula cortex to align the whole-brain EPI to T1 structural image; and 3) Non-linear registration using a combination of FLIRT and FNIRT (Andersson et al., 2007) to align the T1 structural scan to 1mm standard space. Functional MRI scans were resampled once into standard space with a concatenated warp from all three registration steps, and then spatial smoothing in standard space was performed using a Gaussian kernel with 3mm full-width half-maximum using the fslmaths tool.

#### Single-subject general linear model

Single-subject estimates of the general linear model (GLM) were performed using SPM. A GLM was constructed for each of the participants, with a design matrix informed by trial-wise estimates from the RW model of each participant (see above). An additional analysis, using a more classical (non-computational model-based) design matrix, is presented in the Supplementary Material (Supplementary Figures 4 and 6). Alongside the task regressors described below, rigorous de-noising was performed by the inclusion of the following regressors (all described above): the convolved end-tidal CO_2_ regressor plus temporal and dispersion derivatives, six motion regressors trajectories plus their first-order derivatives, physiological noise regressors (provided by the PhysIO toolbox) and ICA components identified as noise.

The regressors of interest in the design matrix were as follows (compare Figure 3A):

1. A ‘Cue’ regressor (80 repeats), with onsets and durations (2.5s) determined by the presentations of visual cues and a magnitude of 1;
2. A ‘Positive prediction’ regressor, with onsets given by the presentation of each corresponding visual cue (when no-resistance was predicted), durations of 0.5s and magnitudes given by 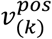 in Equation 5;
3. A ‘Negative prediction’ regressor, with onsets given by the presentation of each corresponding visual cue (when resistance was predicted), durations of 0.5s and magnitudes given by 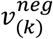 in Equation 6;
4. A ‘No resistance’ stimulus regressor, with onset timings according to the first inspiration that occurred after the presentation of the visual cue, and durations as the remaining time of the potential resistance period (circle in Figure 3), with a magnitude of 1;
5. A ‘Resistance’ stimulus regressor, with onset timings according to the initiation of the inspiratory resistance (identified from the downward inflection of the inspiratory pressure trace) after the presentation of the visual cue, and durations as the remaining time of the resistance period (circle in Figure 3), with a magnitude of 1;
6. A ‘Positive prediction error’ regressor, with onsets given by the start of each corresponding no resistance period, durations of 0.5 s and magnitudes given by 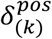 in Equation 7;
7. A ‘Negative prediction error’ regressor, with onsets given by the start of each corresponding resistance period, durations of 0.5 s and magnitudes given by 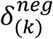 in Equation 8;
8. A ‘Rating period’ noise regressor, with onsets and durations covering the period where participants were asked to rate the difficulty of the previous stimulus, and with a magnitude of 1.

Regressors 1-8 were included in the design matrix after convolution with a standard HRF in SPM12, together with their temporal and dispersion derivatives. Contrasts of interest from this design examined brain activity associated with the average across positive and negative valence for both predictions and prediction errors, as well as the difference due to valence (i.e. positive vs. negative) for both predictions and prediction errors.

#### Group fMRI analysis

Firstly, for the analysis of our entire field of view, contrasts of interest were assessed using random effects group-level GLM analyses based on the summary statistics approach in SPM12. The group-level GLM consisted of a factorial design with both a group mean and group difference regressor. The analyses used a significance level of p<0.05 with family-wise error (FWE) correction at the cluster-level, with a cluster-defining threshold of p<0.001. Secondly, for our region of interest (ROI) analysis, we used FSL’s non-parametric threshold-free cluster enhancement (Smith and Nichols, 2009) within a combined mask of the anterior insula and periaqueductal gray (PAG), as pre-specified in our analysis plan (section 7.5.5). This analysis employed a significance level of p<0.05, with FWE correction across the joint mask. While the anterior insula and PAG have previously been shown to be involved in both conditioned anticipation and perception of inspiratory resistances (Berner et al., 2017; Faull and Pattinson, 2017; Faull et al., 2016, 2018; Paulus et al., 2012; Walter et al., 2020) as well as prediction errors (Roy et al., 2014) and precision (Grahl et al., 2018) towards pain perception, our current analysis considers computational trial-by-trial estimates of interoceptive predictions and prediction errors for the first time. The mask of the anterior insula was taken from the Brainnetome atlas (Fan et al., 2016) (bilateral ventral and dorsal anterior insula regions), and the PAG incorporated an anatomically-defined mask that has been used in previous fMRI publications (Faull and Pattinson, 2017; Faull et al., 2016).

### Multi-modal analysis

#### Multi-modal data

The different task modalities were then combined into a multi-modal analysis to assess both the relationships between and shared variance amongst measures. The data entered into this analysis consisted of:

1. The scores from the four main affective questionnaires that were not used to pre-screen the participants (STAI-S (Spielberger et al., 1970), GAD-7 (Spitzer et al., 2006), ASI-3 (Taylor et al., 2007) and CES-D (Radloff, 1977));
2. The four interoceptive questionnaires (BPQ (Porges, 1995), MAIA (Mehling et al., 2012), PCS-B (Sullivan et al., 1995) and PVQ-B (McCracken, 1997));
3. The four FDT measures (breathing sensitivity, decision bias *c*, metacognitive bias, metacognitive performance Mratio); and
4. The individual peak anterior insula activity associated with both positive and negative predictions, as well as positive and negative prediction errors. Activity was extracted from a 4mm sphere, centred on each participant’s maximal contrast estimate within a Brainnetome atlas mask of the anterior insula (Fan et al., 2016), using the first eigenvariate of the data.

#### Multi-modal correlations and shared variance

A Pearson’s correlation matrix of all 16 included measures was calculated in order to visualise the relationships between all variables. The significance values of the correlation coefficients were taken as p<0.05 (exploratory), and a false discovery rate (FDR) correction for multiple comparisons was applied (using the *mafdr* function in MATLAB). A supplementary non-parametric correlation matrix was additionally calculated using Spearman’s rho values, and these results are presented in Supplementary Table 7.

To assess the shared variance across measures and delineate which measures were most strongly associated with affective qualities, we entered all specified data into a principal component analysis (PCA), following normalisation using z-scoring within each variable. PCA is an orthogonal linear transformation that transforms the *n* × *m* data matrix **X** (participants × measures) into a new matrix **P**, where the dimensions of the variance explained in the data are projected onto the new ‘principal components’ in descending order. Each principal component consists of a vector of coefficients or weights ***W***, corresponding to the contribution of each measure *m* to each component. The PCA also transforms the original *n* × *m* data matrix **X** to map each row (participant) vector *xi* of **X** onto a new vector of principal component scores ***t***_*i*_, given by:

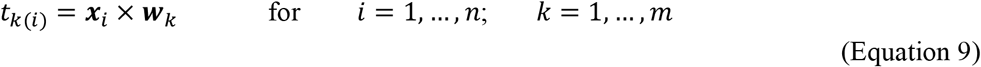

where *t*_k(*i*)_ is the score for each participant *i* within each component *k*. The number of significant components were then determined by comparing the variance explained of each component to a null distribution, created by randomly shuffling (n=1000) the measures from each variable across participants. Components were considered significant if the variance explained was above the 95% confidence interval of the corresponding component’s null distribution.

To assess the relationship between each of the significant components and anxiety, the component scores for low and moderate anxiety were compared using either independent t-tests or Wilcoxon rank sum tests (following Anderson-Darling tests for normality). The significance values of the group differences in component scores were taken as p<0.05 (exploratory), and a false discovery rate (FDR) correction for multiple comparisons (number of significant components) was applied.

An independent code review was performed on all data analysis procedures, and the analysis code is available on GitLab (https://gitlab.ethz.ch/tnu/code/harrison_breathing_anxiety_code).

## Results

### Results overview

Below we present the results from each of our task modalities: questionnaires; a breathing perception/metacognition task (the Filter Detection Task, or FDT); and a novel Breathing Learning Task (BLT), where group-wise comparisons between each of the measures of interest were conducted. The results from the questionnaires and FDT are contextualised by previous findings related to anxiety, while the results from the BLT were validated against an additional unseen dataset and the relationship with anxiety was assessed. We then present the results of a combined multi-modal analysis, where we compared our measures both within and across task modalities. The principal components of these measures were identified to formalise and asses any shared variance between measures, which spanned multiple dimensions of breathing-related interoceptive processing.

### Questionnaire results

The group summaries and comparisons for each of the affective and interoceptive questionnaires (excluding the trait anxiety score that was used for group allocation) are displayed in Figure 1. The group summary values and statistics presented are either mean±standard error (ste) when values were normally distributed and thus compared using unpaired T-tests, or median±inter-quartile range (iqr) when values were not normally distributed and thus compared using Wilcoxon rank sum tests. Scores from all questionnaires of affective symptoms employed were found to be highly significantly different between low and moderate trait anxiety groups: Individuals with moderate levels of trait anxiety demonstrated higher state anxiety (STAI-S mean±ste; low anxiety=25.7±0.7; moderate anxiety=34.1±1.2; t=-6.1; p<0.01), higher levels of anxiety disorder symptoms (GAD-7 median±iqr; low anxiety=1.0±2.0; moderate anxiety=4.0±3.0; Z=-5.9; p<0.01), greater anxiety sensitivity (ASI mean±ste; low anxiety=6.8±0.8; moderate anxiety=18.4±1.5; t=-6.9; p<0.01), and higher levels of depression symptoms (CES-D median±iqr; low anxiety=6.5±3.0; moderate anxiety=14.0±6.0; Z=-6.0; p<0.01).

**Figure 1.**
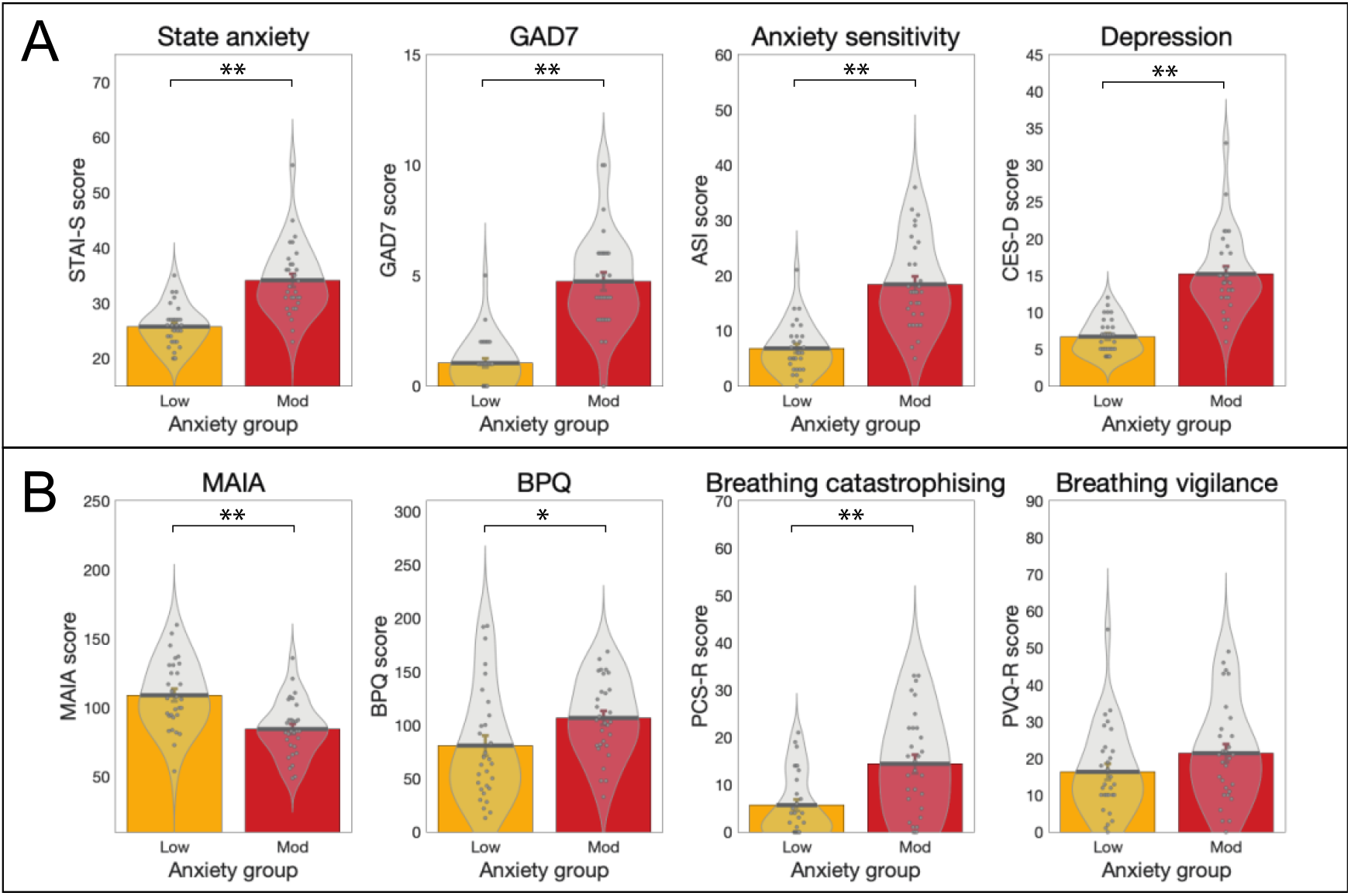
Results from the affective questionnaires (A) and interoceptive questionnaires (B) measured in groups of healthy individuals with either low levels of anxiety (score of 20-25 on the Spielberger Trait Anxiety Inventory, STAI-T) or moderate anxiety (score of 35+ on the STAI-T). Affective questionnaires (A): ‘State anxiety’, Spielberger State Anxiety Inventory; ‘GAD-7’, Generalised Anxiety Disorder Questionnaire; ‘Anxiety sensitivity’, Anxiety Sensitivity Index; ‘Depression’, Centre for Epidemiologic Studies Depression Scale. Interoceptive questionnaires (B): ‘MAIA’, Multidimensional Assessment of Interoceptive Awareness Questionnaire; ‘BPQ’, Body Perception Questionnaire; ‘Breathing catastrophising’, Pain Catastrophising Scale (with the word ‘pain’ substituted for ‘breathing’); ‘Breathing vigilance’, Pain Vigilance Awareness Questionnaire (with the word ‘pain’ substituted for ‘breathing’). *Significant at p<0.05; **Significant following Bonferroni correction for multiple comparisons across all 8 questionnaires. Bar plots represent mean±standard error values, with the distribution of values overlaid in grey. Bar plot code adapted from the CANLAB Toolbox (https://github.com/canlab).

The interoceptive questionnaires we used measured ‘positively-minded’ interoceptive awareness, overall body awareness, breathing symptom catastrophising and breathing symptom vigilance. All except breathing-related vigilance were also found to be significantly different between groups. Individuals with moderate levels of trait anxiety demonstrated reduced ‘positively-minded’ interoceptive awareness (MAIA mean±ste; low anxiety=109.1±4.6; moderate anxiety=84.6±3.7; t=4.2; p<0.01) and greater reports of overall body awareness (BPQ median±iqr; low anxiety=66.0±68.0; moderate anxiety=104.0±52.0; Z=-2.5; p=0.01) in line with previous research (Ewing et al., 2017; Garfinkel et al., 2016b; Mehling, 2016; Paulus and Stein, 2010). Additionally, elevated levels of breathing-related catastrophising were observed in the moderate anxiety group (PCS-B median±iqr; low anxiety=3.5±11.0; moderate anxiety=14.0±17.0; Z=-3.3; p<0.01), while no statistically significant difference was observed for breathing-related vigilance (PVQ-B mean±ste; low anxiety=16.3±2.2; moderate anxiety=21.4±2.5; t=-1.5; p=0.13). Results for sub-component scores and additional questionnaires can be found in Supplementary Figure 1.

### Filter Detection Task results

The group summaries and comparisons for each of the FDT measures are displayed in Figure 2. The FDT output includes the number of filters at perceptual threshold (indicative of perceptual sensitivity, where a greater number of filters indicates lower perceptual sensitivity), decision bias (with *c*<0 indicating a tendency to report the presence of a resistance), metacognitive bias (calculated from average confidence scores) and metacognitive performance (reflecting the congruence between confidence scores and performance accuracy). Individuals with moderate levels of trait anxiety demonstrated both lower perceptual sensitivity (in line with previous findings (Garfinkel et al., 2016a; Tiller et al., 1987)) (filter number median±iqr; low anxiety=3.0±2.0; moderate anxiety=4.0±2.0; Z=-2.4; p=0.01) and lower metacognitive bias (average confidence score median±iqr; low anxiety=6.7±2.2; moderate anxiety=6.2±2.1; Z=2.0; p=0.02) than those with low levels of anxiety, with a similar level of metacognitive performance (Mratio median±iqr; low anxiety=0.8±0.0; moderate anxiety=0.8±0.1; Z=0.7; p=0.23). Decision bias was not found to be different between the groups (decision bias *c* parameter mean±ste; low anxiety=0.15±0.06; moderate anxiety=0.05±0.06; t=1.1; p<0.14) The relationship between greater anxiety and reduced confidence is consistent with results previously observed in the exteroceptive (visual) domain, where decreased confidence related to individual levels of both anxiety and depression (Rouault et al., 2018).

**Figure 2.**
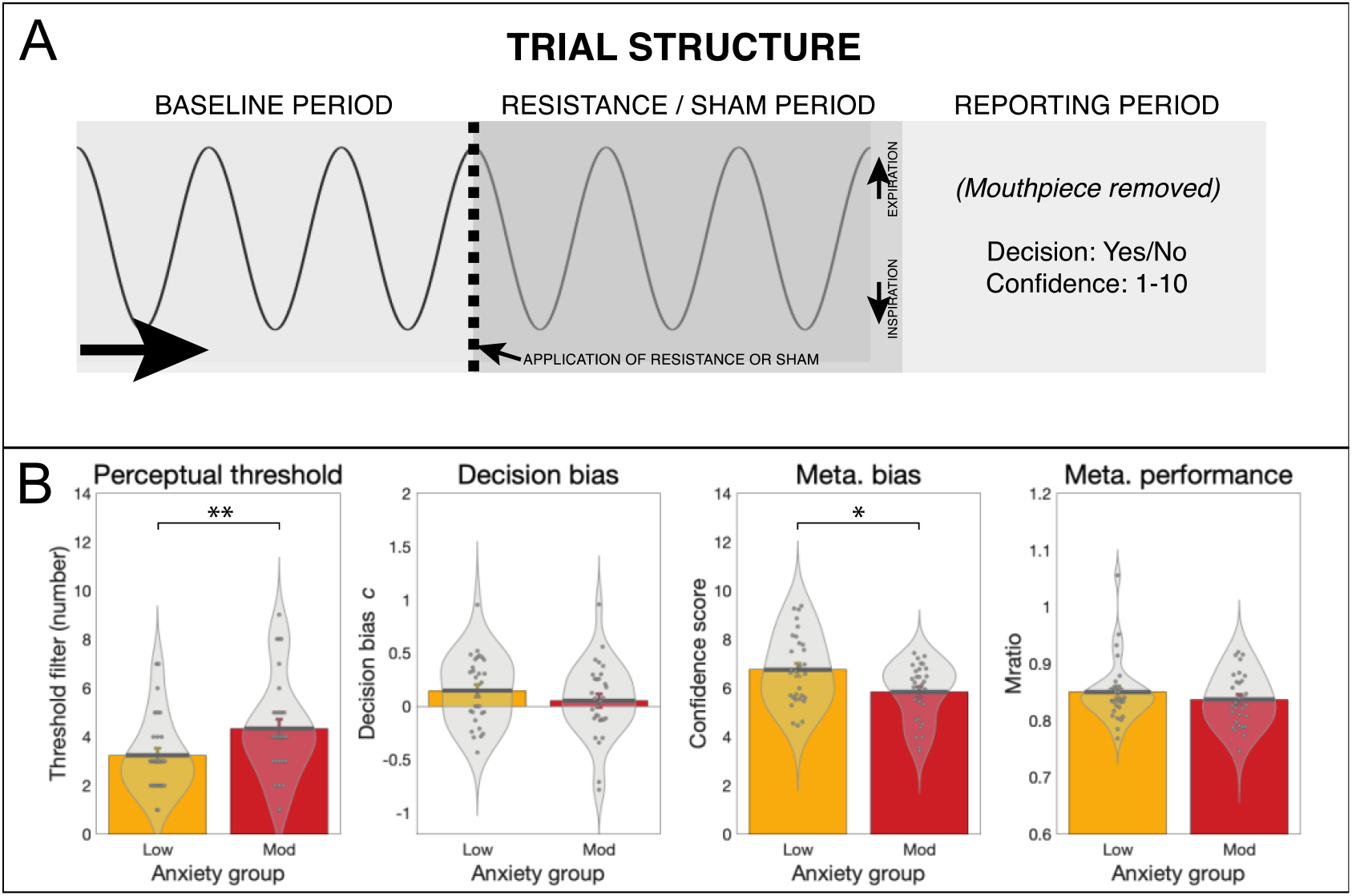
A) Trial structure of the ‘Filter Detection Task’ (FDT). For each trial participants first took three breaths on the system (‘baseline period’), before either an inspiratory resistance or sham was applied. Following three further breaths, participants removed the mouthpiece and reported their decision as to whether a resistance was added (Yes/No), and their confidence in their decision (1-10, 1=not at all confident / guessing; 10=maximally confident in their decision). B) Results from the FDT: Individuals with moderate anxiety (score of 35+ on the STAI-T (Spielberger et al., 1970)) demonstrated a higher (less sensitive) perceptual threshold and lower metacognitive bias (lower average confidence, independent of task accuracy) when compared to individuals with low levels of anxiety (score of 20-25 on the STAI-T (Spielberger et al., 1970)). No difference was found between groups for decision bias (where c values below zero indicate a tendency to report the presence of resistance) nor metacognitive performance (where higher values indicate better metacognitive performance). *Significant at p<0.05; **Significant following Bonferroni correction for multiple comparisons across all FDT measures. Bar plots represent mean±standard error values, with the distribution of values overlaid in grey. Bar plot code adapted from the CANLAB Toolbox (https://github.com/canlab).

### Breathing Learning Task results

#### Behavioural data modelling

When comparing the plausibility of the three alternative models (a Rescorla Wagner, RW; a 2-level Hierarchical Gaussian Filter, HGF2; and a 3-level Hierarchical Gaussian Filter, HGF3) using random effects Bayesian model selection (Rigoux et al., 2014; Stephan et al., 2009), no single model was found to have a protected exceedance probability (PXP) greater than 90% (RW:HGF2:HGF3 PXP=0.30:0.40:0.30, Supplementary Table 4). Therefore, as specified in our analysis plan (https://gitlab.ethz.ch/tnu/analysis-plans/harrison_breathing_anxiety), we conducted our model-based analysis using the conceptually most simple model (the RW model), in accordance with Occam’s Razor. Importantly, the finding that none of the models demonstrated a PXP greater than 90% does not provide any absolute statement about the quality of these models. Rather, this finding indicates that none of the chosen models is conclusively superior to the others in explaining the data. To ensure that the chosen model (RW) provided an adequate explanation of the data, we compared it to a ‘null model’ (i.e. where the choices were due to chance, and not related to any associative learning mechanism) using a likelihood ratio test. We found that in 58 of the 60 participants the RW model fit the behaviour significantly better than the null model, demonstrating that the chosen model captured important aspects of their behaviour. The two participants (one from each anxiety group) who did not show model fits above chance were excluded from any further model-based analyses and comparisons.

To further establish the adequacy of our chosen model to explain learning behaviour in this novel interoceptive learning task, we completed a model validation on 15 additional held-out datasets for the BLT. These participants were not pre-selected for any particular level of anxiety (please see STAR Methods for details). A logistic regression was conducted to assess whether the model prediction trajectory from the original data was able to significantly explain the prediction decisions made by the 15 unseen participants in the validation sample. A representative prediction trajectory from the original 60 participant model fits (the trajectory from the participant with the closest learning rate to the mean) was used in this regression, as well as an intercept term. The beta estimate for the original prediction trajectory was 3.1±0.3 (t=12.1; p=1.0×10^−33^), denoting a highly significant ability of the trajectory to predict unseen data. The beta estimate for the intercept term was -0.2±0.1 (t=-3.1; p=1.7×10^−3^). For a qualitative representation of model fits for both the original and validation data please see Supplementary Figure 3. Both model-based and behavioural parameter comparisons are presented in Table 1. For the estimated model parameters, no difference was observed between the groups for either learning rate (α) or inverse decision temperature (ζ) (Table 1). Results from parameter comparisons between groups including the excluded participants can be found in Supplementary Table 5. For the subjective measures, no difference was observed between the groups for breathing difficulty ratings, while the task-induced anxiety ratings were significantly greater in those with moderate anxiety (Table 1). Additionally, no difference in any physiological measures were observed (Supplementary Table 1), nor in relative head motion during the task (average root mean square displacement (±std) for low anxiety=0.17±0.10 mm; moderate anxiety=0.18±0.07 mm; Wilcoxon rank sum p=0.91).

**Table 1.**
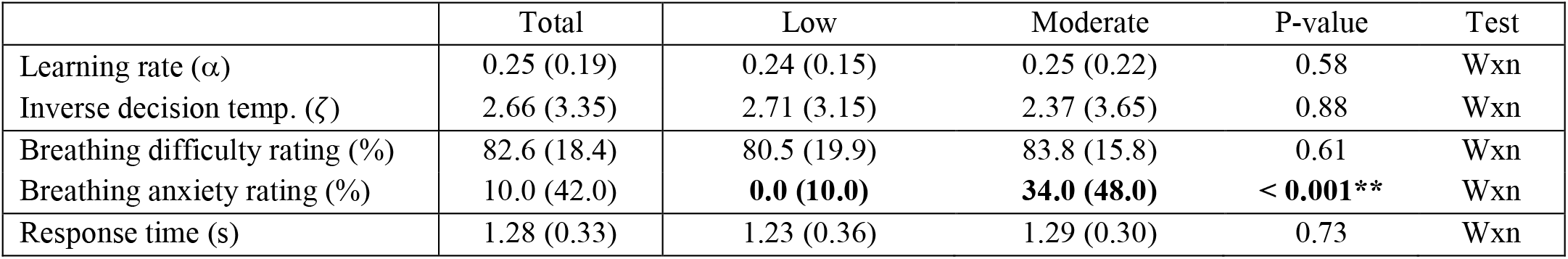
Behavioural and model-based group comparison results from the ‘Breathing Learning Task’ (BLT). All parameters are presented as median±inter-quartile range, and include the model parameter estimates (learning rate, α; inverse decision temperature ζ), the subjective ratings of breathing difficulty (average of the ratings provided following each resistance stimulus) and anxiety (rating provided immediately following the end of the task), and the response times for the predictions made during the task. Abbreviations: Wxn, Wilcoxon rank sum test. If a Wilcoxon rank sum test was utilised, reported values are median ± interquartile range. **Significant difference between groups at p<0.05 with multiple comparison correction for the number of behavioural parameters.

#### Computational modelling of brain activity

The overall and between-group BLT brain activity analysis results are displayed in Figures 4 and 5. In the analysis for the entire field of view, activations related to breathing-related prediction certainty and prediction errors across all participants is shown in Figure 4. Dorsolateral prefrontal cortex (dlPFC), anterior insula (aIns), anterior cingulate cortex (ACC) and middle frontal gyrus (MFG) all demonstrated significant deactivations with overall prediction certainty (i.e. averaged across trials with positive and negative prediction certainty; Figure 4A). In contrast, aIns, ACC, MFG and the periaqueductal gray (PAG) demonstrated significant activations with overall prediction error values (i.e. averaged across trials with positive and negative prediction errors; Figure 4B). A small number of differences due to valence (differences between positive and negative outcomes) were found for prediction errors but not prediction certainty, with negative prediction errors associated with deactivations of left dlPFC and activations of left posterior insula (Figure 4B). While no main effect of anxiety group was observed, an interaction effect was found using the ROI analysis between valence and groups for predictions in the bilateral aIns (Figure 5). In contrast, no group or interaction effects were found for prediction errors. Brain activity associated with inspiratory resistance is provided in Supplementary Figure 6 for comparison with previously published results (Berner et al., 2017; DeVille et al., 2018; Faull and Pattinson, 2017; Faull et al., 2016, 2018; Hayen et al., 2017; Paulus et al., 2012).

**Figure 3.**
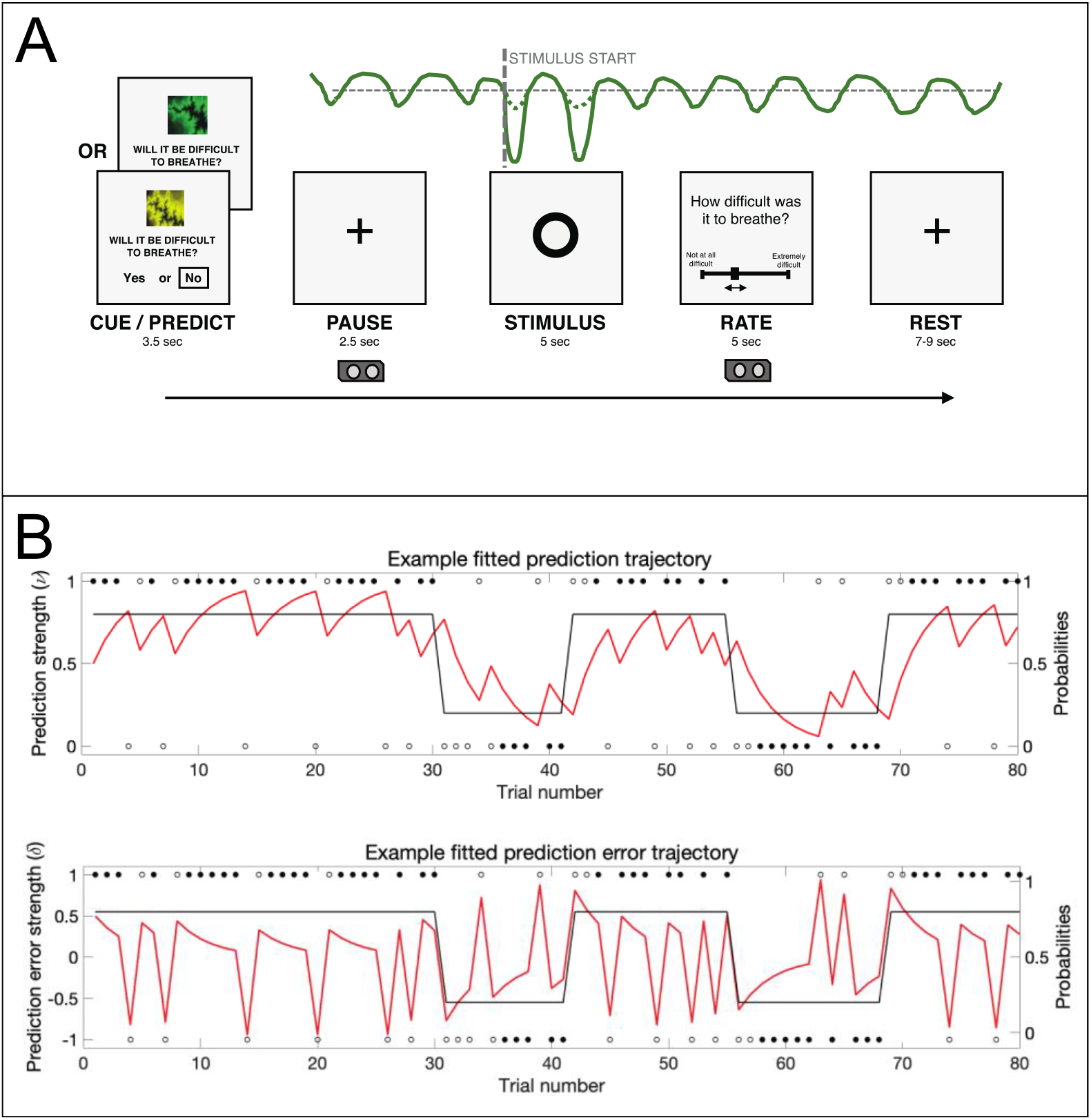
The ‘Breathing Learning Task’ (BLT), used to measure dynamic learning of breathing-related stimuli. A) An overview of the single trial structure, where one of two cues were presented and participants were asked to predict (based on the cue) whether they thought that an inspiratory breathing resistance would follow. When the circle appeared on the screen, either an inspiratory resistance or no resistance was applied for 5 seconds, with the resistance set to 70% of the individual’s maximal inspiratory resistance. After every trial, participants were asked to rate the intensity of the previous stimulus. The trace in green is an example of a pressure trace recorded at the mouth. B) The 80-trial trajectory structure of the probability that one cue predicts inspiratory resistance (black trace), where the alternative cue has an exactly mirrored contingency structure, together with example responses (circles). Filled black circles represent stimuli that were correctly predicted, and open black circles represent stimuli that were not correctly predicted. Example fitted prediction certainty (top) and prediction error (bottom) trajectories are overlaid (red traces). The example trajectories were taken from the participant with the closest learning rate to the mean value across all participants.

**Figure 4.**
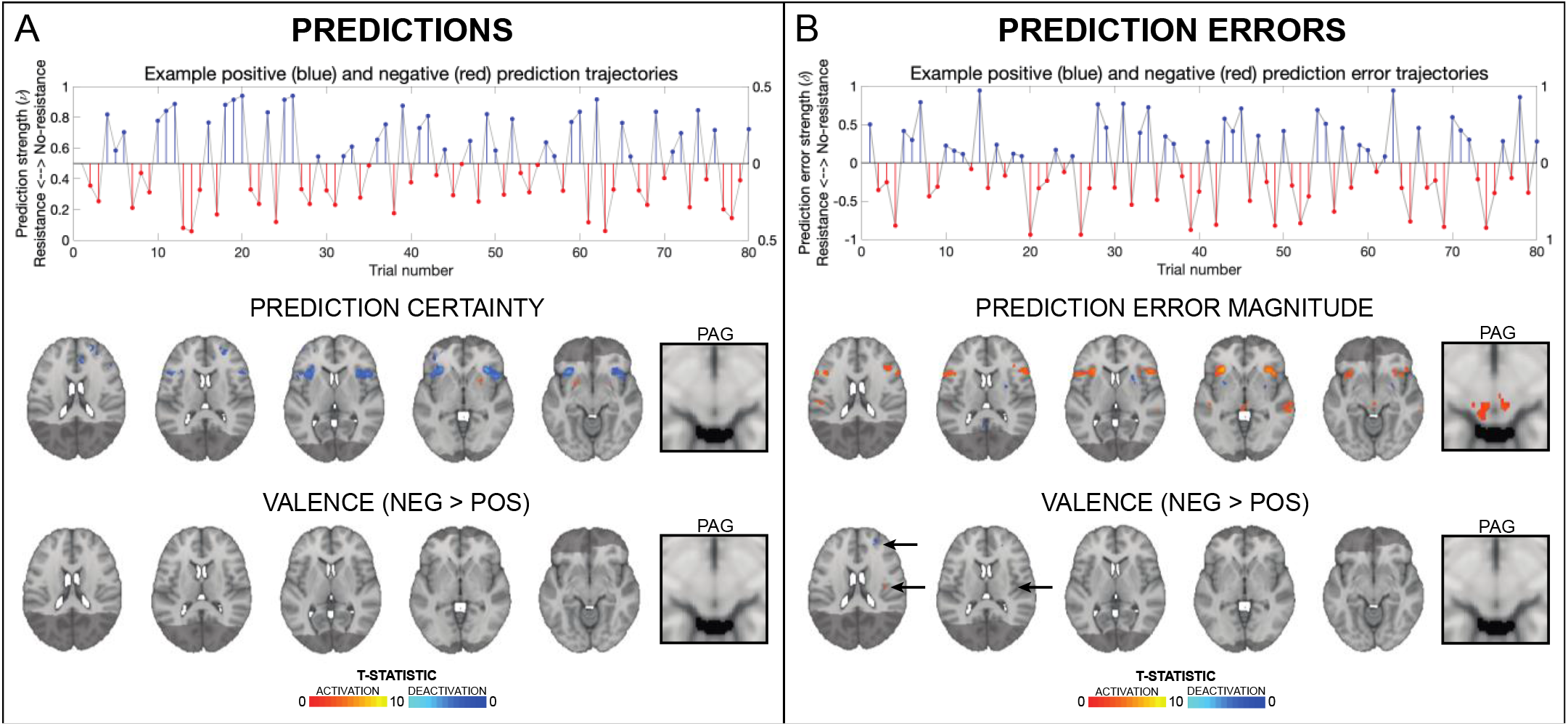
The upper plots demonstrate how estimated prediction (in A) and prediction error (in B) trajectories are encoded as positive (i.e. towards no resistance) and negative (i.e. towards resistance) prediction certainty values and prediction error magnitudes. The example trajectories were taken from the participant with the closest learning rate to the mean value across all participants. The solid grey lines demonstrate the estimated prediction or prediction error traces (in stimulus space). Positive trial values are demonstrated in blue and the negative trial values in red, encoded as distance from 0 (i.e. absolute values; right axes). The brain images represent significant activity across both groups for prediction certainty (averaged over trials with positive and negative prediction certainty) and the influence of valence on prediction certainty (difference between negative and positive predictions), prediction error magnitude (averaged over trials with positive and negative prediction errors) and the influence of valence on prediction error magnitude (difference between negative and positive prediction errors). The images consist of a colour-rendered statistical map superimposed on a standard (MNI 1×1×1mm) brain. The bright grey region represents the coverage of the coronal-oblique functional scan. Significant regions are displayed with a cluster threshold of p<0.05, FWE corrected for multiple comparisons across all voxels included in the functional volume. Abbreviations: PAG, periaqueductal gray.

**Figure 5.**
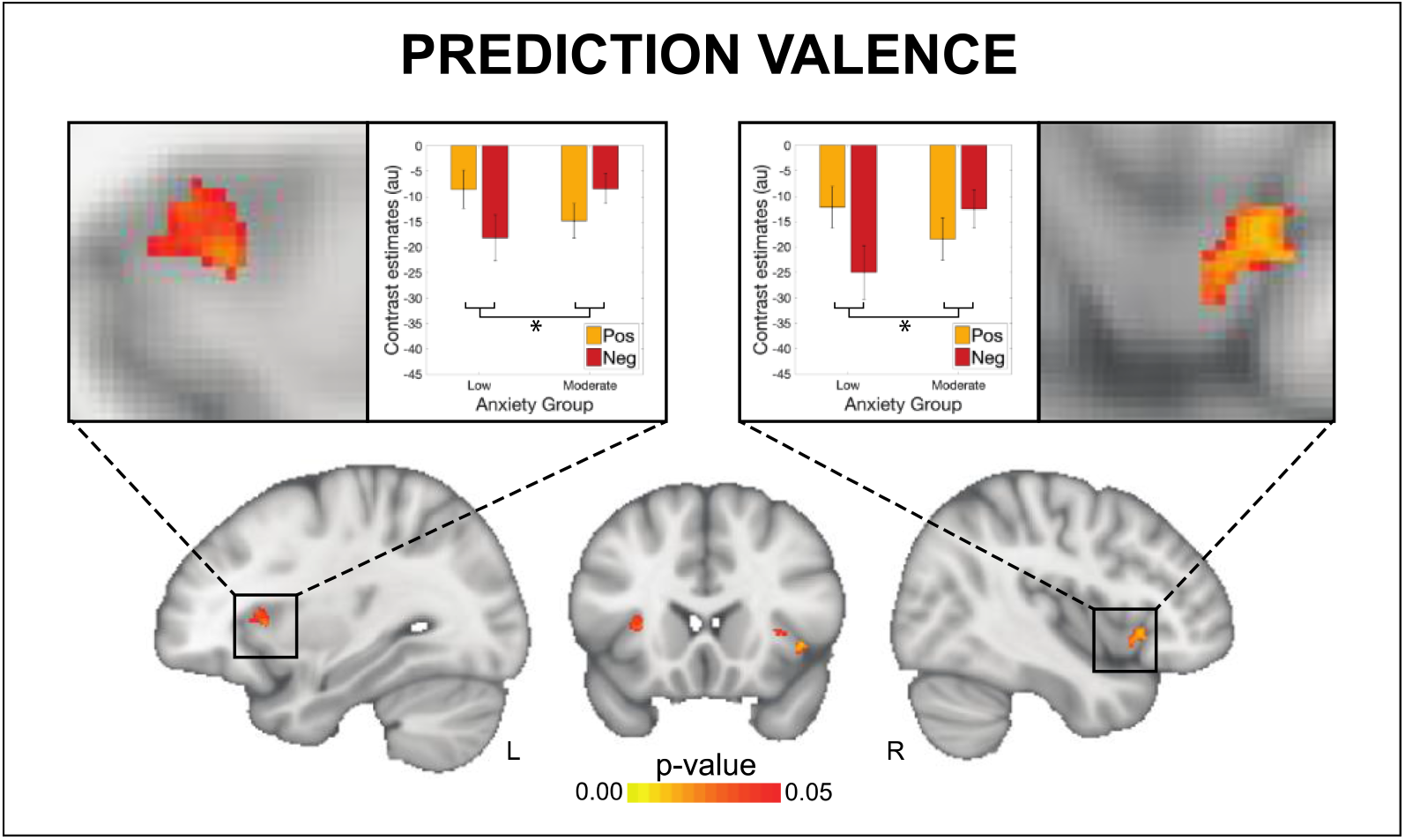
An interaction effect was observed between valence (i.e. trials with positive vs. negative predictions) and anxiety group (low vs. moderate) for activity in the anterior insula related to prediction certainty. The images consist of a colour-rendered statistical map superimposed on a standard (MNI 1×1×1mm) brain. Voxel-wise statistics were performed using non-parametric permutation testing within a mask of the anterior insula and periaqueductal gray, with significant results determined by p<0.05 (corrected for multiple comparisons within the mask).

### Multi-modal analysis results

First, the key measures from each of the different modalities were combined into a multi-modal correlation matrix. This analysis allowed us to assess the relationships both within and across task modalities and across levels of breathing-related interoceptive processing. The full correlation matrix of all 16 included measures is displayed in Figure 6A and Supplementary Table 7. To briefly summarise, the strongest across-task modality correlations were found between all affective and interoceptive questionnaires (Figure 6A). Concerning affective questionnaires and the FDT measures, state anxiety was weakly correlated with the FDT perceptual threshold, decision bias and metacognitive bias, while anxiety sensitivity was additionally weakly related to metacognitive bias. Depression scores were also weakly related to the FDT perceptual threshold. Between the interoceptive and the FDT measures, breathing-related catastrophising was weakly related to metacognitive performance on the FDT. Lastly, between the FDT and aIns activity, metacognitive performance was strongly related to the peak aIns activity associated with negative (i.e. resistance-related) prediction errors, while metacognitive bias was weakly related to aIns activity associated with negative (i.e. resistance-related) prediction certainty. Non-parametric correlations (using Spearman’s rho values) produced highly consistent results, and are presented in Supplementary Table 7.

**Figure 6.**
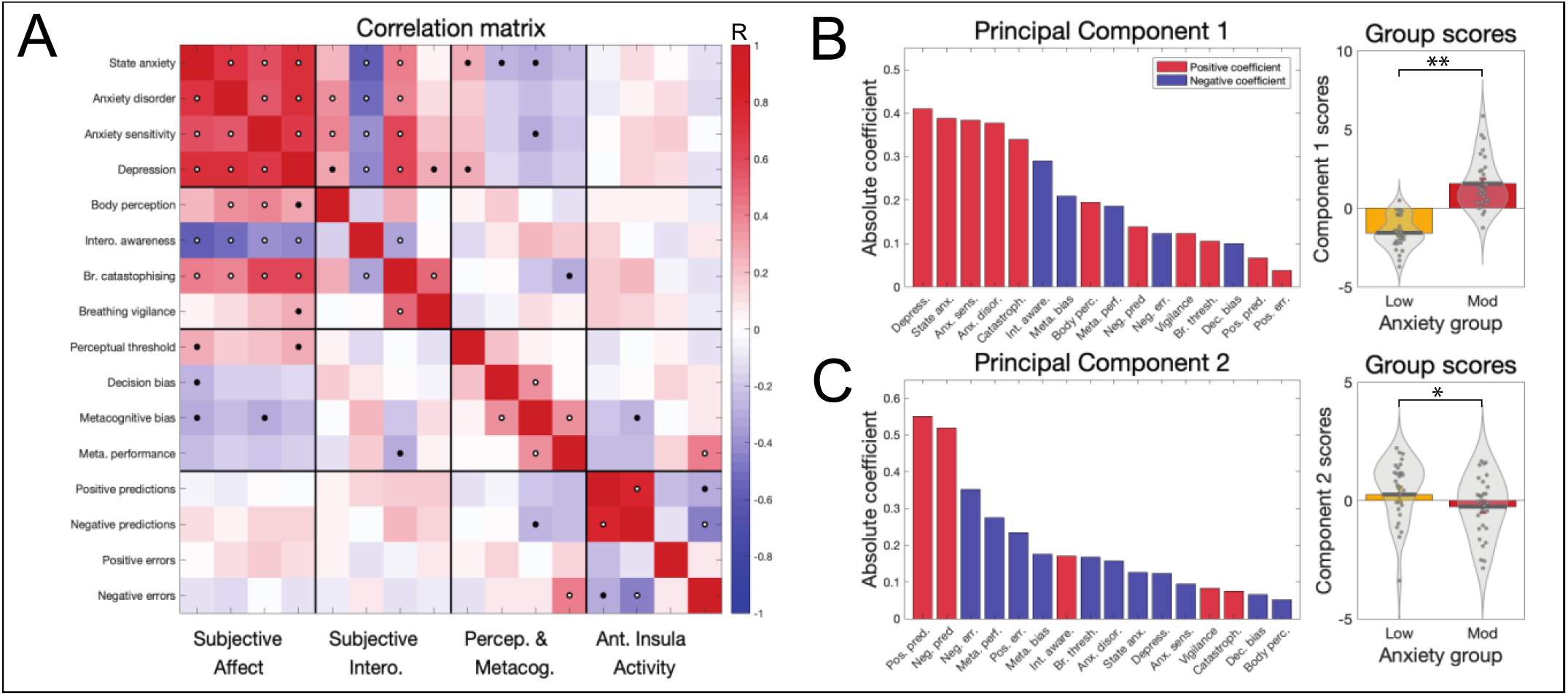
A) Correlation matrix results for the 16 included measures in the multi-modal analysis. Black dots represent significant values at p<0.05, while white dots denote significance with correction for multiple comparisons. B) The weights and group scores of the first significant principal component, where a strong anxiety group difference in component scores is observed. C) The weights and group scores of the second significant principal component, where a weak anxiety group difference in principal component scores is observed. *Significant difference between groups at p<0.05. **Significant difference between groups at p<0.05 with multiple comparison correction for the two significant components. Bar plots (rightmost panels) represent mean±standard error values, with the distribution of values overlaid in grey. Bar plot code adapted from the CANLAB Toolbox (https://github.com/canlab).

#### Principal component analysis

Finally, to assess the extent of shared variance across interoceptive measures, the multimodal data matrix was then subjected to a PCA. This analysis allowed us to delineate how many underlying dimensions may exist within the data, as well as which measures were most strongly associated with trait anxiety. Two principal components (PC) were found to be significant, where the variance explained with each component was above the 95% confidence interval of its null distribution. The properties of each of these significant components are displayed in Figure 6B and 6C. The first PC demonstrated a highly significant (p<1×10^−11^) difference in scores between the anxiety groups. Correspondingly, the greatest coefficients within the first PC were from the affective measures of depression scores, state anxiety, anxiety sensitivity and anxiety disorder scores. Additionally, breathing-related catastrophising and negative interoceptive awareness also had strong coefficient values, followed by negative metacognitive bias (i.e. lower confidence scores), body perception scores (from the BPQ) and negative metacognitive performance (i.e. lower metacognitive performance). In contrast, the second PC demonstrated a weak difference (p=0.05) in scores between the anxiety groups. This component had the highest coefficient scores from the peak aIns activity related to positive and negative prediction certainty, as well as negative coefficients for negative prediction errors, metacognitive performance and positive prediction errors.

## Discussion

### Main findings

Interoceptive abilities are thought to be tightly linked to affective properties such as anxiety. Here we have provided a unifying analysis by characterising this relationship across multiple interoceptive levels in the breathing domain, including novel findings of altered brain activity within the aIns when processing dynamic breathing predictions. This study is the first to demonstrate brain activity related to dynamic interoceptive learning, and, specifically, activity in the insula that is related to breathing-related prediction error and prediction certainty in a trial-by-trial fashion. Notably, aIns activity related to the certainty of predictions about breathing resistance was found to differ between trait anxiety levels. Furthermore, this study is also amongst the first to simultaneously tackle multiple levels of interoceptive processing, using breathing as a salient and accessible channel of body perception. The tasks employed reflected the broad range of targeted processes; not only were questionnaires employed that spanned affect and body perceptions, but behavioural data from two different tasks were assessed by separate computational models. These analyses allowed for formal assessments of both breathing-related interoceptive learning and metacognition, including the first computational assessment of trial-by-trial learning in the interoceptive domain as well as applying state-of-the-art models of metacognition to interoception of breathing. Our multi-modal approach revealed that not only is the relationship between breathing-related interoception and trait anxiety broad, it is most strongly detected (i.e. greatest PCA weights: Figure 6) at the higher levels of interoceptive processing, which includes specific subjective measures of interoceptive beliefs (often termed ‘interoceptive sensibility’ (Garfinkel et al., 2015)) followed by metacognitive aspects of breathing perceptions. Notably, the peak aIns brain activity associated with breathing-related interoceptive learning appeared to be largely independent of other interoceptive measures, with the exception of negative prediction error-related brain activity and metacognitive performance.

### Affect and levels of breathing-related interoception

Beyond consequences at single levels of interoceptive processing, here we aimed to assess how the relationship with trait anxiety may cross multiple interoceptive levels related to breathing. Employing PCA (with permutation testing) allowed us to identify any components that share common variance within our multi-modal dataset, and additionally assess the relative contribution of our measures to each dimension (Figure 6B). Here we found that all affective qualities loaded strongly onto the first principal component, with the greatest additional contributions from subjective measures of negative interoceptive awareness and breathing-related catastrophising. General body awareness and the breathing-related metacognitive measures (bias and performance) were the next largest contributors to this shared variance, followed by the perceptual sensitivity and decision bias parameters, and lastly peak aIns activity from the BLT. These results suggest that the relationship with anxiety is particularly prominent at the level of subjective interoceptive beliefs in the breathing domain, which are thought to exist in the higher levels of interoceptive space (Critchley and Garfinkel, 2017), followed by metacognitive insight into breathing perception. In comparison, the relationship of trait anxiety to lower-level properties such as interoceptive sensitivity (Critchley and Garfinkel, 2017; Garfinkel et al., 2015, 2016b, 2016a) appear to be present but less prominent in the breathing domain. However, it must be noted that quantifying these higher interoceptive levels may be less noisy in comparison to measuring psychophysical properties such as sensitivity, and thus the relationship with anxiety might be most easily detected rather than being inherently stronger.

Although strong relationships were observed between affective qualities and many of our interoceptive breathing measures, a sparse number of correlations were found between interoceptive measures themselves, and in particular across task modalities (Figure 6A). These findings support the idea that there are potentially separable levels of breathing-related interoception as proposed (Critchley and Garfinkel, 2017). The only notably strong cross-modal relationship was found between metacognitive performance and aIns activity, where greater insight into breathing sensitivity correlated with greater aIns activity during negative prediction errors. This relationship may reflect a previously proposed contribution of error-processing towards metacognitive awareness, where deviations between actual and predicted bodily inputs are propagated to metacognitive areas via interoceptive brain structures such as the aIns (Stephan et al., 2016).

### Novel measures of dynamic interoceptive predictions and prediction errors

Many theories surrounding anxiety have hypothesised an important role of altered predictions regarding upcoming threat (Bach, 2015; Mogg et al., 2000; Simmons et al., 2006), and in particular interoceptive threat (Paulus and Stein, 2010; Paulus et al., 2019) in the anterior insula (Allen, 2020; Bossaerts, 2010; Carlson et al., 2010; Paulus and Stein, 2006; Tan et al., 2018). While numerous studies have employed inspiratory resistive loads to evoke threatening interoceptive stimuli (Alius et al., 2013; Berner et al., 2017; Faull and Pattinson, 2017; Faull et al., 2016, 2018; Hayen et al., 2017; Leupoldt and Dahme, 2005; Leupoldt et al., 2009; Paulus et al., 2012; Stoeckel et al., 2016; Walter et al., 2020), the BLT approach presented here is, to the authors’ knowledge, the first investigation of dynamic (trial-by-trial) brain activity associated with model-based interoceptive predictions and prediction errors for respiration. By fitting an associative learning model to each participant’s behavioural responses, we could quantify both the certainty of predictions and magnitude of prediction errors on each trial. We could then compare both the parameter estimates and the brain activity associated with these computational quantities, with a particular focus on the aIns and PAG (Allen, 2020; Grahl et al., 2018; Paulus and Stein, 2006; Roy et al., 2014; Singer et al., 2009) (Figure 4). Here, we observed evidence for a relationship between anxiety and aIns reactivity to threat valence in the prediction domain (Figure 5). Specifically, while individuals with low trait anxiety demonstrated greater aIns deactivation that scaled with predictions of breathing resistance compared to no resistance, the opposite was true in individuals with moderate trait anxiety (creating an interaction effect). This demonstrates a shift in the aIns processing of threat valence with different levels of anxiety, in line with hypothesised differences in brain prediction processing (Paulus and Stein, 2006, 2010; Paulus et al., 2019). In comparison, no anxiety group differences or interactions were found in the breathing prediction error domain, contrasting with some previously proposed hypotheses (Barrett and Simmons, 2015; Brewer et al., 2021; Paulus and Stein, 2006, 2010).

Beyond the anterior insula and independent of anxiety, multiple (and largely consistent) proposals have been made, inspired by predictive coding and related theories of brain function, regarding which brain networks might be involved in processing interoceptive predictions and prediction errors (Allen, 2020; Barrett, 2016; Barrett and Simmons, 2015; Craig, 2009; Khalsa et al., 2017; Kleckner et al., 2017; Manjaly and Iglesias, 2020; Marlow et al., 2019; Owens et al., 2018; Paulus et al., 2019; Pezzulo et al., 2015, 2018; Quadt et al., 2018; Seth, 2013; Smith et al., 2017; Stephan et al., 2016). These proposed networks are loosely hierarchical in structure and typically assign metacognitive processes to higher cortical areas (e.g. prefrontal cortex; PFC) while interoceptive predictions are thought to originate from regions that may serve as an interface between interoceptive and visceromotor function (e.g. aIns and anterior cingulate cortex; ACC). In these concepts, prediction errors have two different roles: on the one hand, they are thought to be sent up the cortical hierarchy of interoceptive regions in order to update predictions in aIns and ACC (Barrett and Simmons, 2015; Pezzulo et al., 2015; Seth et al., 2012); on the other hand, they are thought to determine regulatory signals, sent from visceromotor regions and brainstem structures like the PAG to the autonomic nervous system and bodily organs (Petzschner et al., 2017; Stephan et al., 2016).

While these theories have received considerable attention, there has been little empirical evidence thus far. In particular, we are not aware of any studies that have demonstrated, using a concrete computational model, trial-by-trial prediction and prediction error activity in interoceptive areas. Here, we report evidence of relevant computational quantities being reflected by activity within several areas of a putative interoceptive breathing network. While activity related to trial-wise prediction certainty was found in higher structures such as dorsolateral PFC, ACC and aIns, prominent prediction error responses were not only found in aIns and ACC, but also in the midbrain PAG (Figure 4). Importantly, concerning predictions, widespread brain activity was found to be mainly related to prediction *un*certainty, where BOLD activity was decreased in proportion to increases in the certainty of predictions (Feldman and Friston, 2010; Friston, 2005). Furthermore, it is notable that the anterior insula displayed both deactivation for more certain predictions and activation for greater prediction errors. This might reflect the proposed critical role of aIns in the representation and updating of models of bodily states (Allen, 2020; Bergh et al., 2017; Manjaly and Iglesias, 2020; Paulus and Stein, 2006; Paulus et al., 2019; Seth, 2013; Stephan et al., 2016; Walter et al., 2020), given that greater certainty (precision of beliefs) reduces and greater prediction errors increase belief (model) updating (Petzschner et al., 2017).

Our PAG findings are of particular interest. While the PAG has been previously noted during anticipation of certain breathing resistance stimuli (Faull and Pattinson, 2017; Faull et al., 2016) and has been related to the precision of prior beliefs about placebo-induced reductions in pain intensity (Grahl et al., 2018), here we observed that PAG activity did not appear to be related to the extent of prediction certainty towards upcoming breathing stimuli (Figure 4). Concerning prediction error activity in the PAG, this has previously been demonstrated in relation to pain (Roy et al., 2014); here, we found PAG activity in relation to the magnitudes of trial-wise interoceptive prediction errors (Figure 4B), consistent with a role of PAG in homeostatic control (Stephan et al., 2016).

Finally, overall prediction and prediction error-related activity did not appear to be dissociated between anterior and posterior insula cortices (respectively), as has been previously hypothesised (Allen, 2020; Barrett and Simmons, 2015; Stephan et al., 2016). However, a small valence difference in prediction errors was observed, with negative prediction errors (the unexpected presence of inspiratory resistance stimuli) producing greater activity in the right posterior insula than positive prediction errors (the unexpected absence of inspiratory resistance stimuli; Figure 4). It is therefore possible that the representation of homeostatically relevant inputs in the posterior insula is enhanced for events that may negatively impact homeostasis. However, these results are the first demonstration of brain activity related to dynamic interoceptive prediction and prediction errors; furthermore, the functional images from this study do not have the resolution required for layer-specific identification of prediction and prediction error processing in the insula. Additionally, the represented prediction error-related activity may be specific to the breathing domain within interoceptive processing. This latter caveat of course also applies to the wider results presented here; as only one interoceptive channel (i.e. inspiratory resistances within the breathing domain) was tested, we cannot assume these results would translate to other interoceptive processes, e.g. related to cardiac or gastric states.

## Conclusions

The relationship between anxiety and breathing crosses multiple levels of the interoceptive hierarchy. In particular, anxiety and associated affective dimensions appear to be most strongly related to subjective negative body awareness and catastrophising about breathing symptoms, followed by metacognitive measures related to breathing perception. Furthermore, a novel interaction between trait anxiety group and valence was found within the aIns associated with dynamic prediction certainty (but not prediction errors) of breathing-related interoceptive processing. More generally, this study provides the first empirical demonstration of brain activity associated with dynamic (trial-by-trial) interoceptive learning. These results provide new and comprehensive insights into how anxiety is related to levels of interoceptive processing in the breathing domain, and provide evidence of brain activity associated with trial-wise predictions and prediction errors about bodily states in interoceptive breathing networks.

## Author contributions

OKH, FP and KES designed the study; OKH, LN, SM, RL, FH and KB acquired the data; OKH and SJH analyzed the data with input from KES, AJH, SF, SI and FV; OKH and KES wrote the manuscript, with edits from all remaining authors.

## Acknowledgements

The authors would like to thank Professor Klaas Pruessmann and Dr Lars Kasper for their work establishing and supporting the MRI protocol. OKH (née Faull) was supported by a Marie Skłodowska-Curie Postdoctoral Fellowship from the European Union’s Horizon 2020 research and innovation programme under the Grant Agreement No 793580. SF was supported by the UZH Forschungskredit Postdoc FK-18-046, as well as the ETH Zurich Postdoctoral Fellowship Program and the Marie Curie Actions for People COFUND program FEL-49 15-2. FV was supported by the Fondation Deniker, the Fondation pour la Recherche Médicale, and the Fondation Bettencourt Schueller. SJH was supported by the grant #2017-403 of the Strategic Focus Area “Personalized Health and Related Technologies (PHRT)” of the ETH Domain. KES was supported by the René and Susanne Braginsky Foundation and the University of Zurich.

## Supplementary Material

**Supplementary Figure 1:**
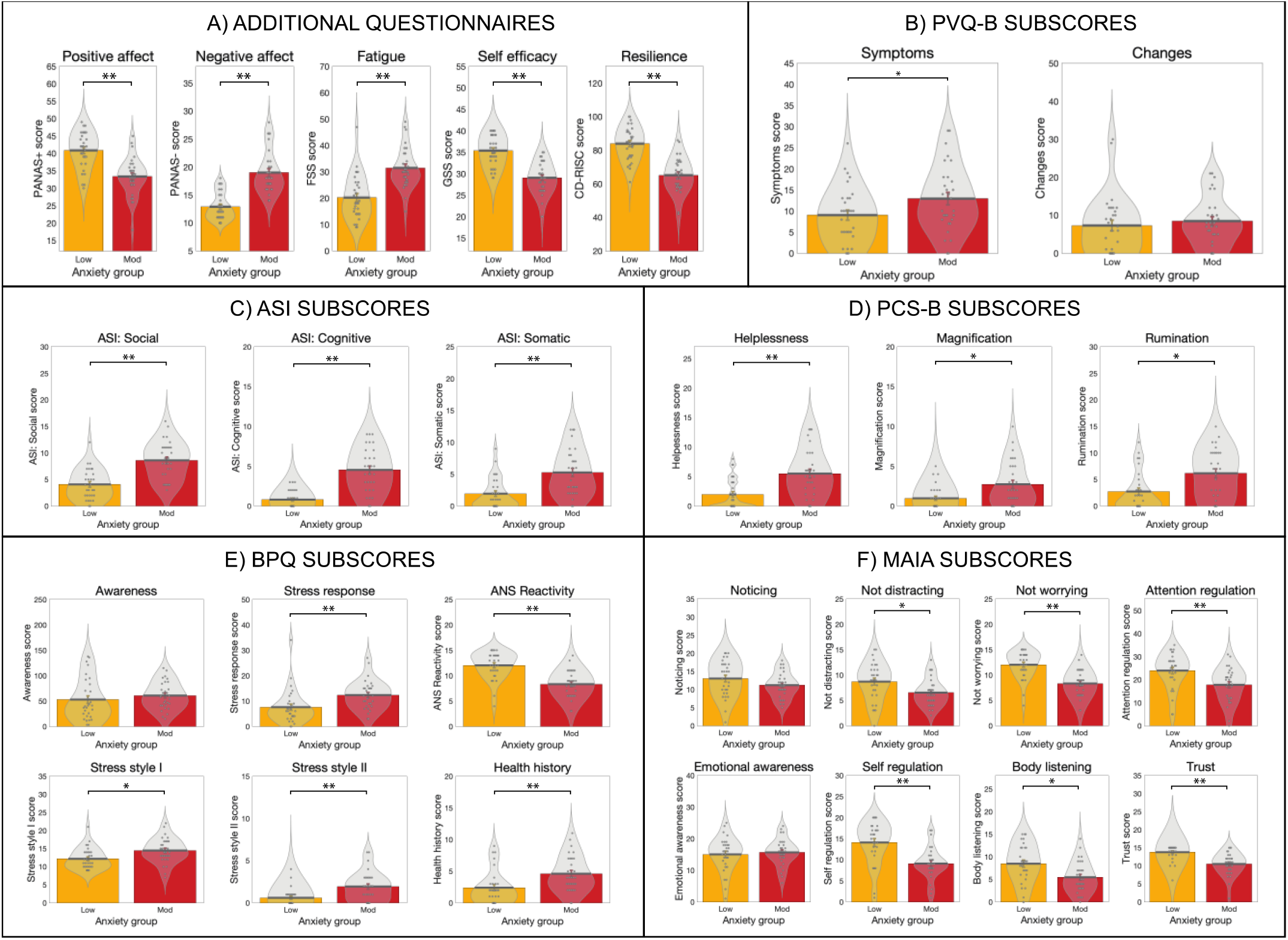
Related to Figure 1. A) Results from the additional questionnaires measured in groups of healthy individuals with either low levels of anxiety (score of 20-25 on the Spielberger Trait Anxiety Inventory, STAI-T) or moderate anxiety (score of 35+ on the STAI-T). Questionnaires: ‘Positive affect’ and ‘Negative affect’ from the Positive Affect Negative Affect Schedule (PANAS-T), ‘Fatigue’ from the Fatigue Severity Scale (FSS), ‘Self efficacy’ from the General Self-Efficacy Scale, and ‘Resilience’ from the Connor-Davidson Resilience Scale. B) Results from the sub-scores of the Pain Vigilance Awareness Questionnaire (with the word ‘pain’ substituted for ‘breathing’, PVQ-B). C) Results from the sub-scores of the Anxiety Sensitivity Index (ASI-3) questionnaire. D) Results from the sub-scores of the Pain Catastrophising Scale (with the word ‘pain’ substituted for ‘breathing’, PCS-B). E) Results from the sub-scores of the Body Perception Questionnaire (BPQ). F) Results from the sub-scores of the Multidimensional Assessment of Interoceptive Awareness Questionnaire (MAIA). Bar plots represent mean±standard error values, with the distribution of values overlaid in grey. **Significant at p<0.01, with no correction for multiple comparisons (exploratory results). Bar plots represent mean±standard error values, with the distribution of values overlaid in grey. Bar plot code adapted from the CANLAB Toolbox (https://github.com/canlab).

**Supplementary Figure 2:**
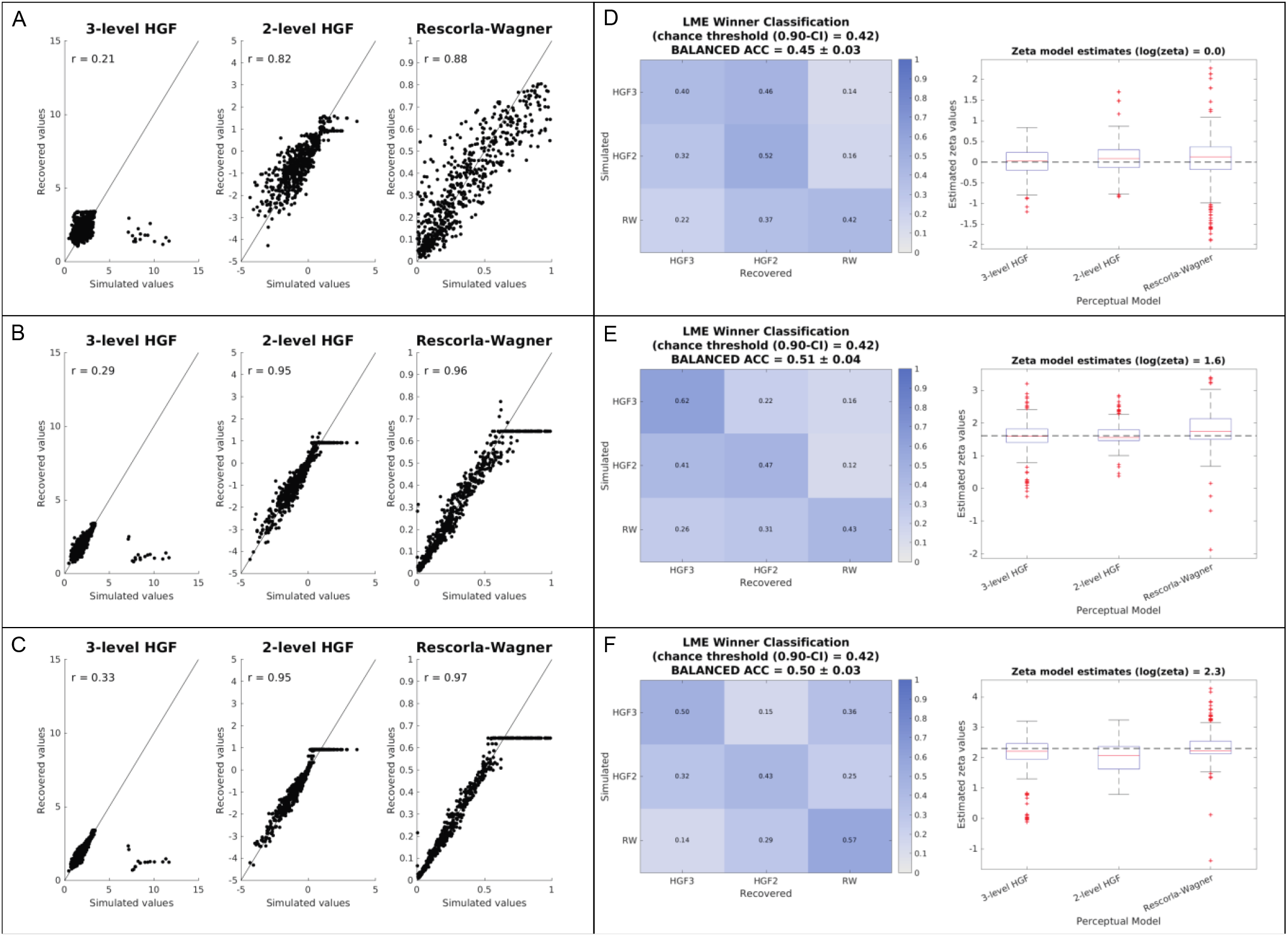
Related to Figure 3. A-C) Demonstration of parameter recovery using simulated participants from the prior distributions presented in Supplementary Table 2. 60 simulated participant responses were generated using 10 different seed values, totalling n=600 simulations plotted here. D-E) Demonstration of model identifiability using simulated participants from the prior distributions presented in Supplementary Table 2. 60 simulated participant responses were generated using 10 different seed values, and the confusion matrices, balanced accuracy and zeta estimates are the average values across the 10 simulation runs. Three noise levels were used for the simulations, with an inverse decision temperature (ζ) of 1 (A,D), 5 (B,E) and 10 (C,E), representing very noisy (ζ = 1) to very deterministic (ζ = 10) settings.

**Supplementary Figure 3:**
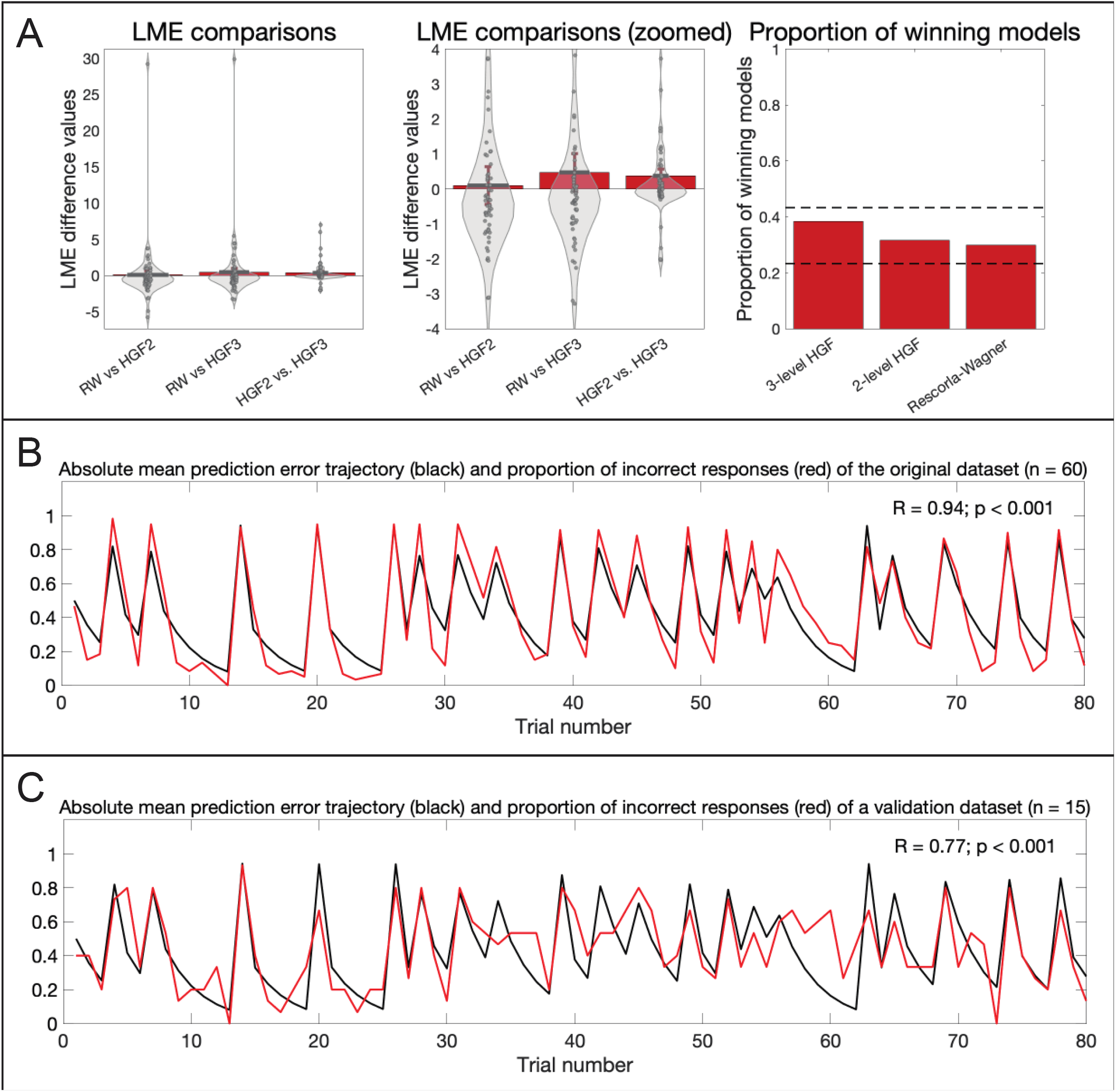
Related to Figure 3. A) Model comparison: Estimated log model evidence (LME) values across all participants. Left and middle: Comparisons between each model pair, where bar plots represent mean±standard error values of the specified differences in LME, with the distribution of values overlaid in grey. Bar plot code adapted from the CANLAB Toolbox (https://github.com/canlab). Right: Proportion of highest LME values across all 60 participants. Dotted lines represent upper and lower 95% confidence intervals for chance, and all proportions of winner classifications lie within the chance range. B) Model validation in original data: Comparison of the mean prediction error trajectory (black) against the proportion of participants giving incorrect responses at each trial (red) for the original dataset (n = 60). The close correlation between these trajectories demonstrates the extent to which the chosen model (the Rescorla Wagner) captures important aspects of participant performance. C) Model validation in unseen data: Comparison of the mean prediction error trajectory from the original dataset (black) against the proportion of participants giving incorrect responses at each trial (red) for a validation dataset (n = 15), who were not preselected for any specific anxiety level. The correlation between these trajectories demonstrates the extent to which the average model fit for the original data is able to capture important aspects of participant performance on **unseen** data. The prediction error trace in (B) and (C) is the absolute prediction error trajectory from the participant with the closest learning rate to the mean across all participant model fits. The prediction error trace is represented in stimulus space (see Supplementary Figure 4 for transformation procedures).

**Supplementary Figure 4:**
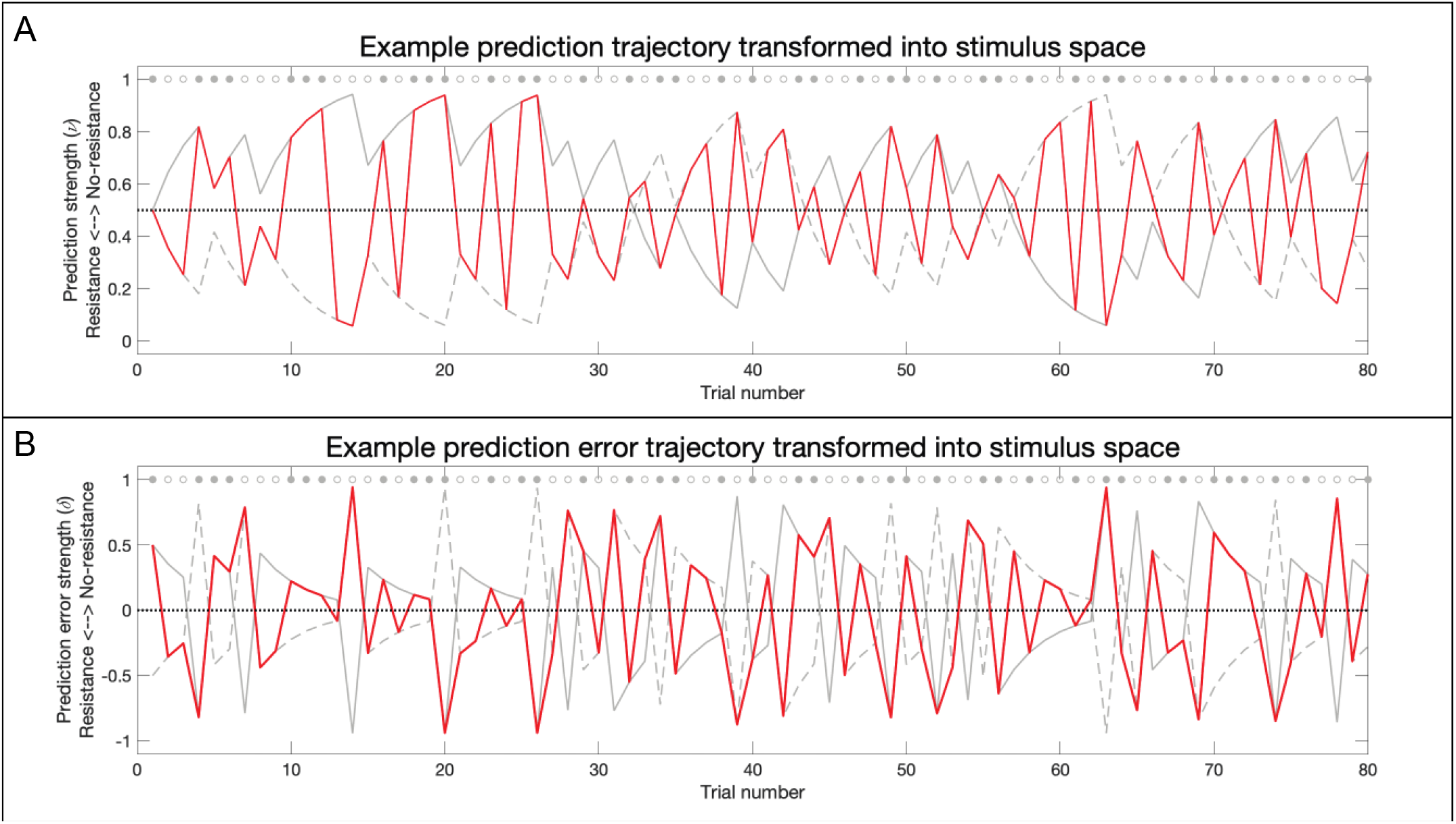
Related to Figures 4 and 5. Transformation of the prediction (A) and prediction error (B) trajectories from contingency space to stimulus space. In Panel A, the fitted trajectory (in contingency space) is demonstrated by the solid grey line, where the value 1 is assigned when one cue (cue 1) predicts no resistance and the opposing cue (cue 2) predicts a resistance (the value 0 is assigned for the opposing conditions). The trajectories were then transformed into stimulus space, where a value of 1 was assigned when no resistance was delivered, while a value of 0 was assigned when a resistance was delivered. For this transformation, a mirrored trajectory was firstly generated (dashed grey line) to represent the second cue, as the participants were explicitly told that the cues acted as a pair that had opposite probabilities (20% or 80%) of predicting resistance. The solid grey trajectory thus represents the cue that started with an 80% probability of being followed by no resistance in stimulus space (cue 1), while the dashed grey line represents the cue that started with a 20% probability of being followed by no resistance (cue 2). The values at each trial were taken from the trajectory of the cue that was presented at that trial: either cue 1 (trials with a closed grey circle) or cue 2 (trials with an open grey circle). The same transformation was performed on the prediction error trajectories in Panel B, where the solid grey line represents the prediction error associated with cue 1, while the dashed grey line represents the prediction error associated with cue 2. The example trajectories were taken from the participant with the closest learning rate to the mean value across all participants.

**Supplementary Figure 5:**
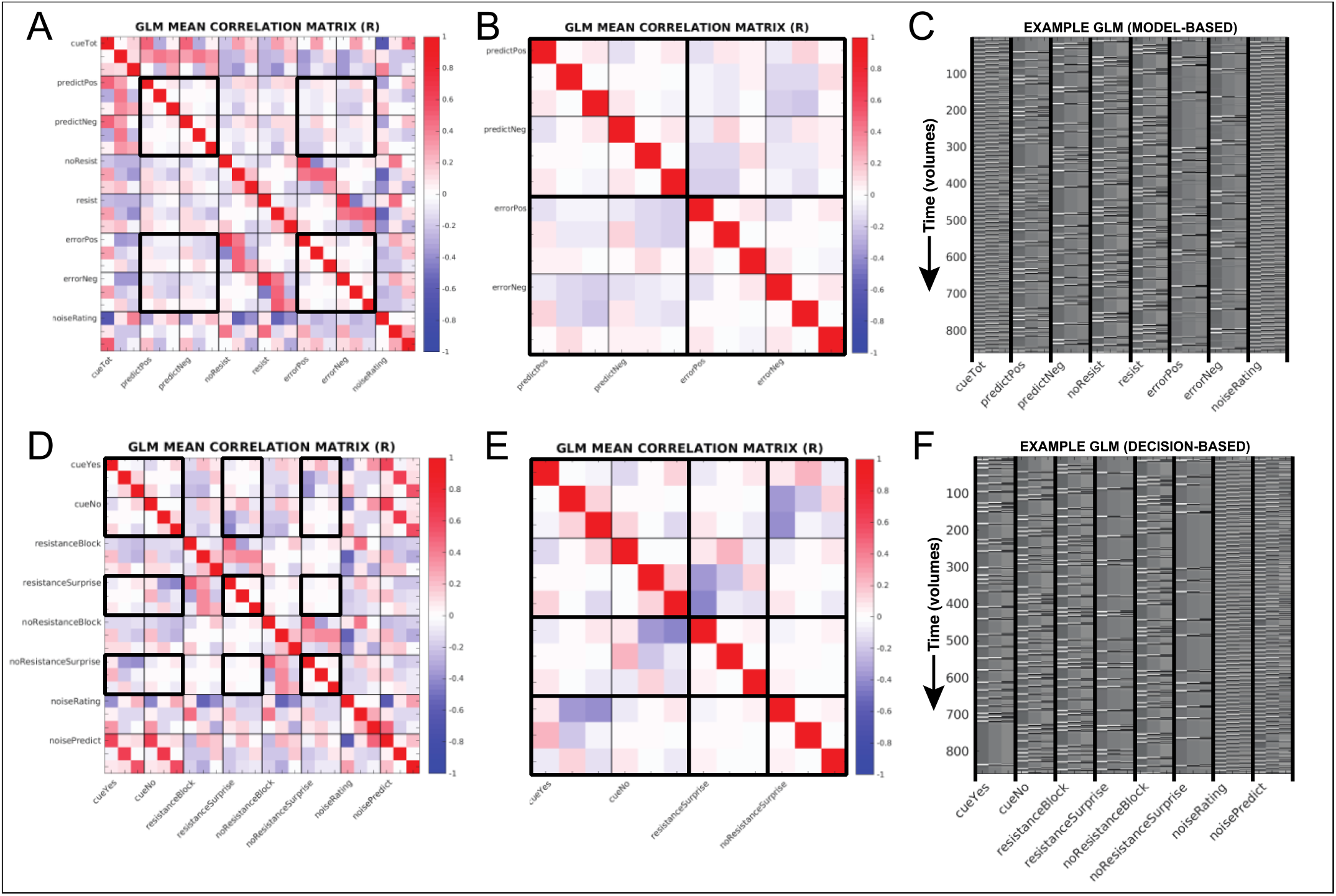
Related to Figure 4 and 5. A and D) Average correlation matrices (using Fischer’s R-to-Z transformation prior to averaging) from all single subject general linear models used in the model-based (A) and decicion-based (D) fMRI analyses (noise regressors not shown). B and E) Targeted correlation matrices to demonstrate the relationship between predictions and errors for A and D, respectively. C) An example general linear model from a single participant model-based fMRI analysis. Each of the main regressors also include a temporal and dispersion derivative. Please see STAR methods for a full description of each regressor. F) An example general linear model from a single participant decision-based fMRI analysis. Each of the main regressors also include a temporal and dispersion derivative, and consist of: 1) ‘cueYes’ – the time periods covering the presentation of cues where the participant predicted an upcoming resistance; 2) ‘cueNo’ – the time periods covering the presentation of cues where the participant predicted no upcoming resistance; 3) ‘resistanceBlock’ – the stimulus periods when resistance was applied, from the onset of the inspiration against the increased inspiratory pressure following the presentation of the circle cue to the end of the circle presentation; 4) ‘resistanceSurprise’ – resistance stimuli that were surprising (i.e. when participant had predicted no resistance), with an onset at the beginning of the corresponding stimulus period and a duration of 0.5 seconds; 5) ‘noResistanceBlock’ – the stimulus periods when no resistance was applied, from the onset of the first inspiration following the presentation of the circle cue to the end of the circle presentation; 6) ‘noResistanceSurprise’ – no-resistance stimuli that were surprising (i.e. when participant had predicted resistance), with an onset at the beginning of the corresponding stimulus period and a duration of 0.5 seconds; 7) ‘noiseRating’ – the periods at the end of each trial where the participant rated the intensity of the previous stimulus, with an onset at the beginning and duration that encompassed the length of the rating period. 8) ‘noisePredict’ – the button press periods during the cue presentation, with an onset given by the response time for the button press on each trial and a duration of 0.5 seconds. Noise regressors not shown: the convolved end-tidal carbon dioxide trace (plus temporal and dispersion derivatives), the RETROICOR and convolved respiratory volume per unit of time (RVT) regressors provided by the PhysIO toolbox, 6 motion regressors and 6 extended motion regressors (derivatives), and the timeseries of all identified noise components from the independent component analysis conducted during preprocessing were also included in the model.

**Supplementary Figure 6:**
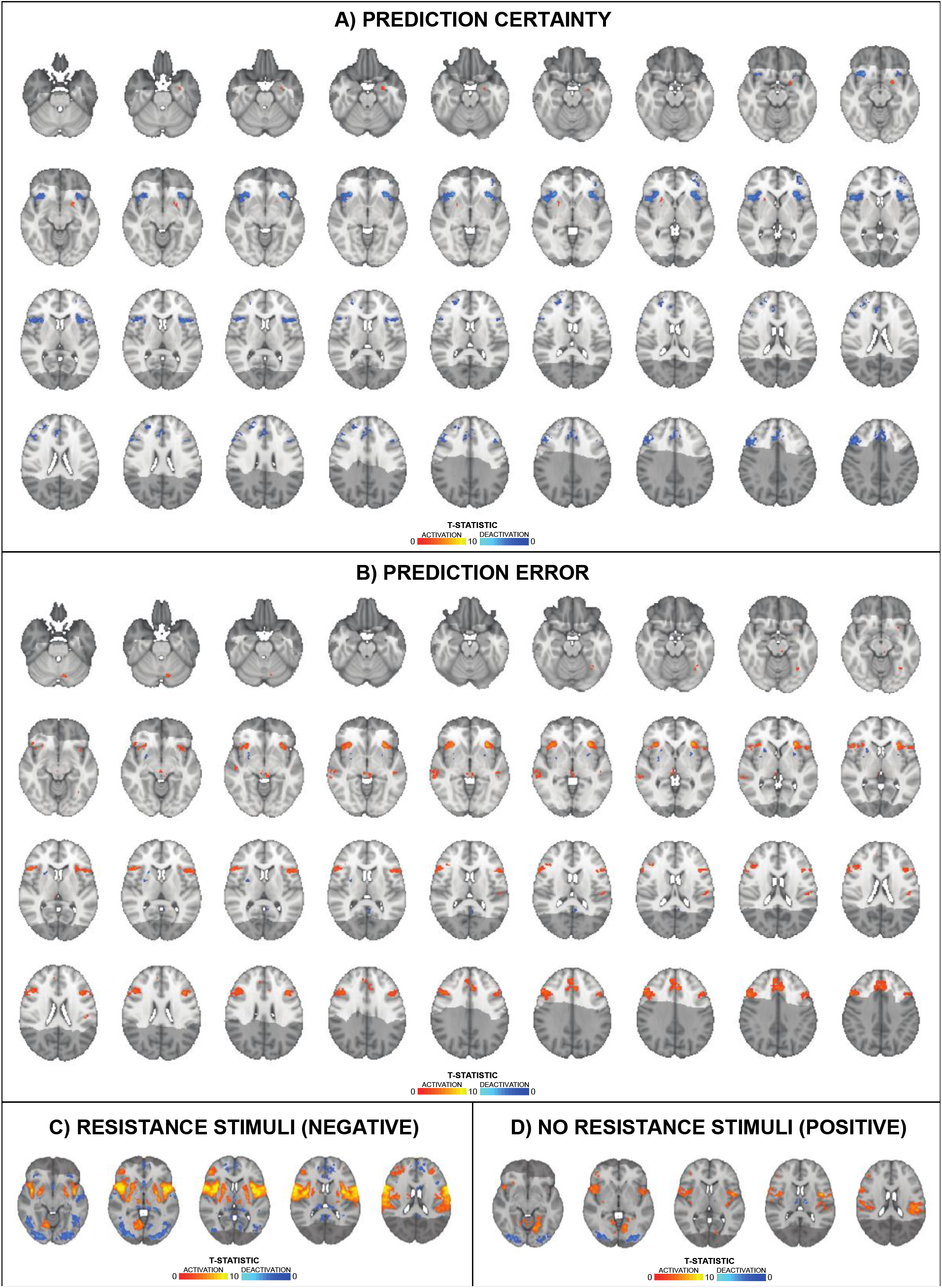
Related to Figures 4 and 5. A) Significant BOLD activity associated with prediction certainty, averaged over trials with positive and negative prediction certainty. B) Significant BOLD activity associated with prediction error magnitude, averaged over trials with positive and negative prediction errors. C) Significant BOLD activity associated with inspiratory resistance periods. D) Significant BOLD activity associated with no resistance periods. and no inspiratory resistance (bottom panel) stimulus periods. The images consist of a colour-rendered statistical map superimposed on a standard (MNI 1×1×1mm) brain. The bright grey region represents the coverage of the coronal-oblique functional scan. Significant regions are displayed with a cluster threshold of p<0.05, FWE corrected for multiple comparisons across all voxels included in the functional volume. Images in A and B are an expanded view of those presented in Figure 4.

**Supplementary Figure 7:**
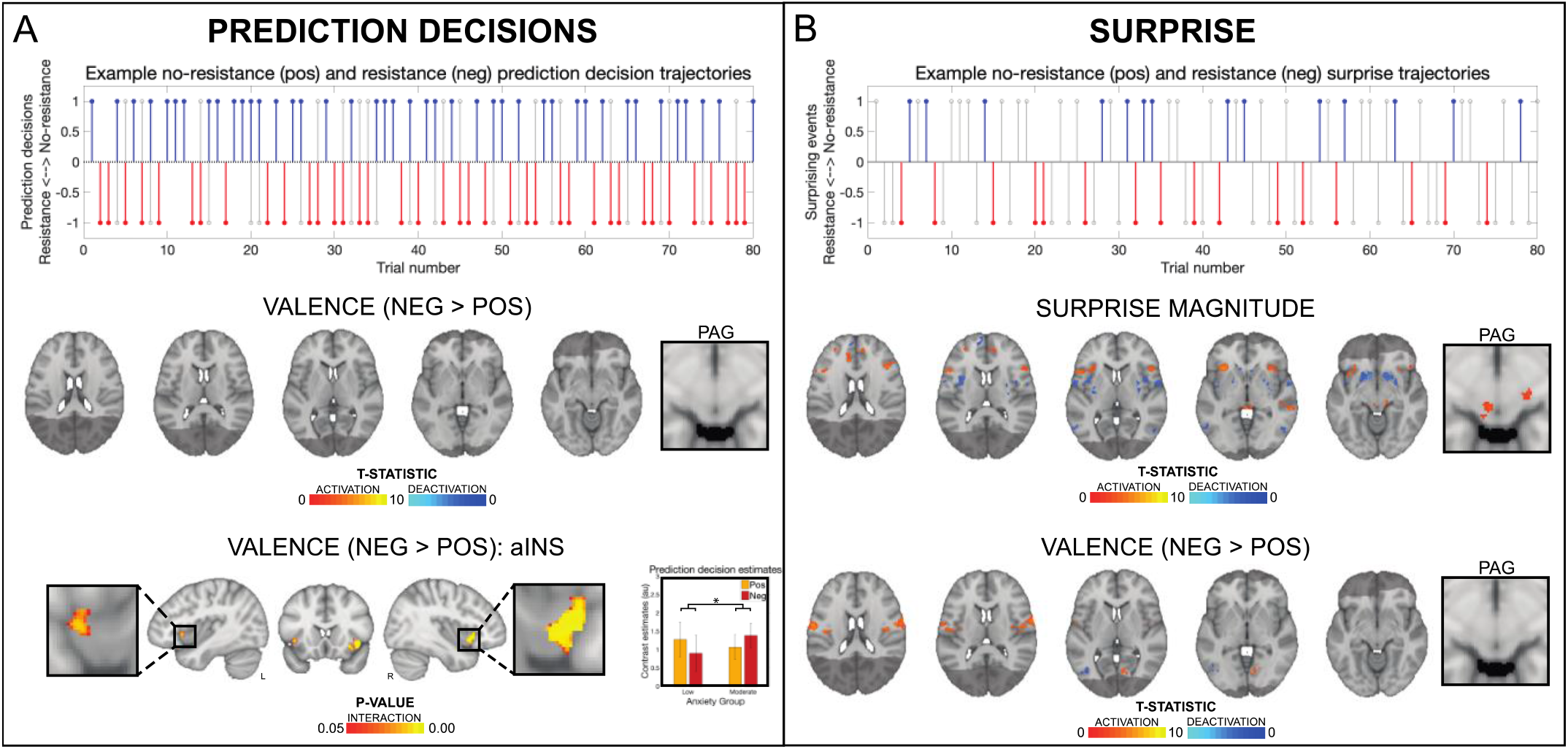
Related to Figures 4 and 5. Overall results from the non-computational decision-based analyses of the ‘Breathing Learning Task’ (BLT). The plots in both (A) and (B) demonstrate how prediction decisions (in A) and surprise (in B) trajectories are encoded into positive (i.e. towards no resistance) and negative (i.e. towards resistance, red) values. The grey lines in both plots represent the stimulus at each trial, while the blue (positive) and red (negative) lines in (A) denote the prediction decisions (prior to the stimulus) and in (B) the surprising events (where the prediction decision was incorrect). In both trajectories the dotted black line denotes the boundaries between positive and negative valence, and the distance from the dotted line is taken as the final value (i.e. absolute values). The brain images in (A) represent the influence of valence on prediction decisions (difference between negative and positive decisions), while there is no equivalent representation of overall predictions in a binary decision model compared to the computational model design (the average over positive and negative prediction decisions simply represents the cue presentation). The bottom panel of brain images in (A) demonstrate an interaction effect between valence (i.e. positive vs. negative) and anxiety group (low vs. moderate) for the anterior insula activity related to the valence of the prediction decisions of interoceptive breathing stimuli. Here, voxel-wise statistics were performed using non-parametric permutation testing within a mask of the anterior insula and periaqueductal gray, with significant results determined by p<0.05 (corrected for multiple comparisons within the mask). The brain images in (B) represent the activity associated with average surprise (average over positive and negative surprise trajectories) and the influence of valence on surprise (difference between negative and positive surprise trajectories). The images consist of a colour-rendered statistical map superimposed on a standard (MNI 1×1×1mm) brain. The bright grey region represents the coverage of the coronal-oblique functional scan. Significant regions are displayed with a cluster threshold of p<0.05, FWE corrected for multiple comparisons across all voxels included in the functional volume. Abbreviations: PAG, periaqueductal gray

**Supplementary Table 1:** Related to Figure 3. Physiological summaries and group comparison results from the stimulus periods of the ‘Breathing Learning Task’ (BLT). Abbreviations: Wxn, Wilcoxon rank sum test; Ttest, students independent T-test. If a Wilcoxon rank sum test was utilised, reported values are median ± interquartile range.

|  | Total | Low | Moderate | P-value | Test |
| --- | --- | --- | --- | --- | --- |
| **RESISTANCE** |  |  |  |  |  |
| 70% of maximum inspiratory pressure | -58.8 (24.5) | -63.4 (23.8) | -56.0 (28.0) | 0.52 | Wxn |
| Avg measured pressure (cmH<sub>2</sub>O) | -4.0 (3.7) | -4.2 (3.8) | -3.8 (2.9) | 0.72 | Wxn |
| Max measured pressure (cmH<sub>2</sub>O) | -7.3 (4.9) | -7.3 (6.7) | -7.3 (3.9) | 0.68 | Wxn |
| Avg breathing rate (bpm) | 13.6 (6.7) | 14.6 (5.6) | 13.0 (10.1) | 0.39 | Wxn |
| Avg breathing depth (%) | 103.9 (25.5) | 97.0 (23.7) | 105.9 (26.1) | 0.29 | Ttest |
| Heart rate (bpm) | 66.3 (13.5) | 66.9 (12.5) | 66.0 (15.6) | 0.69 | Ttest |
| **NO RESISTANCE** |  |  |  |  |  |
| Avg measured pressure (cmH<sub>2</sub>O) | -0.1 (0.1) | -0.1 (0.1) | -0.1 (0.1) | 0.05 | Wxn |
| Max measured pressure (cmH<sub>2</sub>O) | -0.8 (0.5) | -0.8 (0.8) | -0.7 (0.4) | 0.59 | Wxn |
| Avg breathing rate (bpm) | 14.8 (5.1) | 14.9 (5.0) | 14.7 (5.8) | 0.92 | Ttest |
| Avg breathing depth (%) | 106.6 (17.0) | 105.0 (17.0) | 108.8 (21.7) | 0.35 | Ttest |
| Heart rate (bpm) | 66.7 (12.3) | 67.6 (12.1) | 66.6 (14.0) | 0.84 | Wxn |

**Supplementary Table 2:** Related to Figure 3. Parameter configurations and priors for each of the candidate models. If the prior variance is set to 0 for a parameter then it is not estimated, and the prior mean and variance for the estimated parameters (in bold) were taken from the maximum likelihood fits of the pilot participant data. Prior means are given in native space, prior variances in estimation (transformed) space.

| *Rescorla Wagner* |  |  |  |
| --- | --- | --- | --- |
| Parameter | Prior mean | Prior variance | Transformation |
| $v^{(0)}$ | 0.5 | 0 | logit |
| $\alpha$ | **0.29** | **2.54** | **logit** |
| Observation model $\zeta$ | **2.14** | **3.33** | **log** |
| *Hierarchical Gaussian Filter (2-level)* |  |  |  |
| Parameter | Prior mean | Prior variance | Transformation |
| $\mu_2^{(0)}$ | 0 | 0 | none |
| $\mu_3^{(0)}$ | 1 | 0 | none |
| $\sigma_2^{(0)}$ | 0.1 | 0 | log |
| $\sigma_3^{(0)}$ | 1 | 0 | log |
| $\rho_2$ | 0 | 0 | none |
| $\rho_3$ | 0 | 0 | none |
| $\kappa_1$ | 1 | 0 | log |
| $\kappa_2$ | 0 | 0 | log |
| $\omega_2$ | **-0.70** | **1.61** | **none** |
| $\omega_3$ | $-\infty$ | 0 | none |
| Observation model $\zeta$ | **2.14** | **1.17** | **log** |
| *Hierarchical Gaussian Filter (3-level)* |  |  |  |
| Parameter | Prior mean | Prior variance | Transformation |
| $\mu_2^{(0)}$ | 0 | 0 | none |
| $\mu_3^{(0)}$ | 1 | 0 | none |
| $\sigma_2^{(0)}$ | 0.1 | 0 | log |
| $\sigma_3^{(0)}$ | 1 | 0 | log |
| $\rho_2$ | 0 | 0 | none |
| $\rho_3$ | 0 | 0 | none |
| $\kappa_1$ | 1 | 0 | log |
| $\kappa_2$ | **2.39** | **0.30** | **log** |
| $\omega_2$ | -3 | 0 | none |
| $\omega_3$ | -6 | 0 | none |
| Observation model $\zeta$ | **2.14** | **1.21** | **log** |

**Supplementary Table 3:** Related to Figure 3. Parameter recovery metrics for each of the candidate models. R values are Pearson Correlation Coefficients that have been Fisher’s Z-transformed prior to averaging across the 10 simulation runs, and then back-transformed into R values. Each simulation run consisted of 60 simulated synthetic datasets sampled from the prior distribution of values for the perceptual model, repeated at each noise level (ζ = [1,5,10]), and recovered using MAP estimation.

| Rescorla Wagner |  |  |  |  |  |  |
| --- | --- | --- | --- | --- | --- | --- |
| | $\zeta = 1$ | | $\zeta = 5$ | | $\zeta = 10$ | |
|  | Mean | Std | Mean | Std | Mean | Std |
| $\alpha$ R | 0.88 | 0.07 | 0.96 | 0.14 | 0.97 | 0.11 |
| $\zeta_{est}$ | 1.08 | 1.70 | 5.82 | 1.56 | 10.62 | 1.61 |
| Hierarchical Gaussian Filter (2-level) |  |  |  |  |  |  |
| | $\zeta = 1$ | | $\zeta = 5$ | | $\zeta = 10$ | |
|  | Mean | Std | Mean | Std | Mean | Std |
| $\omega_2$ R | 0.83 | 0.10 | 0.95 | 0.14 | 0.96 | 0.15 |
| $\zeta_{est}$ | 1.11 | 1.47 | 5.05 | 1.36 | 7.89 | 1.57 |
| Hierarchical Gaussian Filter (3-level) |  |  |  |  |  |  |
| | $\zeta = 1$ | | $\zeta = 5$ | | $\zeta = 10$ | |
|  | Mean | Std | Mean | Std | Mean | Std |
| $\kappa_2$ R | 0.32 | 0.31 | 0.49 | 0.49 | 0.56 | 0.59 |
| $\zeta_{est}$ | 1.01 | 1.35 | 4.92 | 1.50 | 8.52 | 1.75 |

**Supplementary Table 4:**
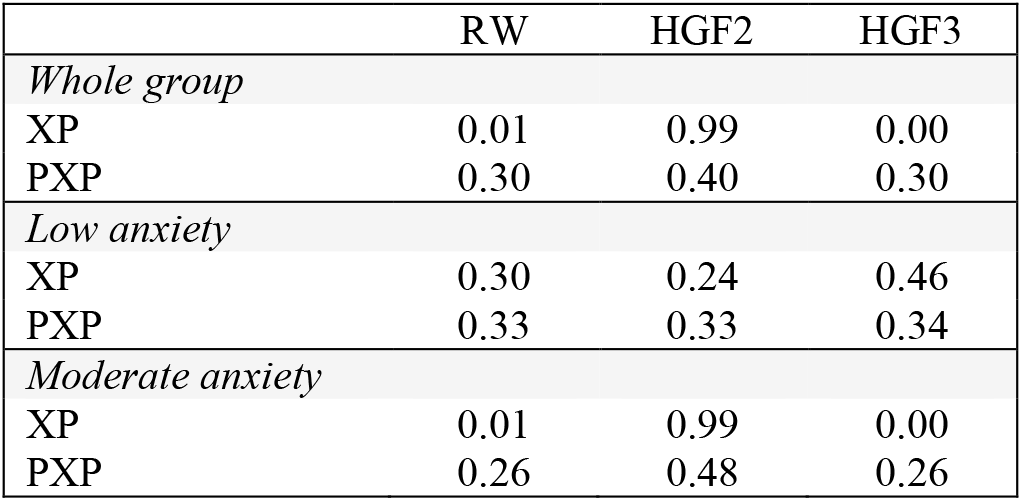
Related to Figure 3. Whole and individual-group model comparison results. Abbreviations: XP, exceedance probability; PXP, protected exceedance probability.

|  | RW | HGF2 | HGF3 |
| --- | --- | --- | --- |
| *Whole group* |  |  |  |
| XP | 0.01 | 0.99 | 0.00 |
| PXP | 0.30 | 0.40 | 0.30 |
| *Low anxiety* |  |  |  |
| XP | 0.30 | 0.24 | 0.46 |
| PXP | 0.33 | 0.33 | 0.34 |
| *Moderate anxiety* |  |  |  |
| XP | 0.01 | 0.99 | 0.00 |
| PXP | 0.26 | 0.48 | 0.26 |

**Supplementary Table 5:**
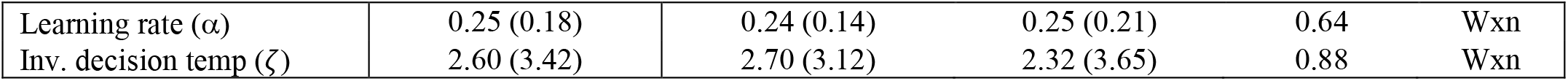
Related to Table 1. Behavioural group comparison results from the ‘Breathing Learning Task’ (BLT), with fitted perceptual and response model parameters (learning rate, α; and inverse decision temperature ζ). All participants included in the comparison. Abbreviations: Wxn, Wilcoxon rank sum test; Ttest, students independent T-test. If a Wilcoxon rank sum test was utilised, reported values are median ± interquartile range.

|  |  |  |  |  |  |
| --- | --- | --- | --- | --- | --- |
| Learning rate ($\alpha$) | 0.25 (0.18) | 0.24 (0.14) | 0.25 (0.21) | 0.64 | Wxn |
| Inv. decision temp ($\zeta$) | 2.60 (3.42) | 2.70 (3.12) | 2.32 (3.65) | 0.88 | Wxn |

**Supplementary Table 6:** Related to Figure 3. Exploratory regression analysis conducted on the fitted model learning rate parameter and the subjective ratings of breathing difficulty and anxiety. Regression parameters consisted of trait anxiety scores (from the STAI-T questionnaire), depression scores (from the CES-D questionnaire) and gender (male=1). **Significant coefficient at p<0.05 with multiple comparison correction for the three exploratory regression models.

|  | Trait anxiety | p-value | Depression score | p-value | Gender (male) | p-value |
| --- | --- | --- | --- | --- | --- | --- |
| Learning rate | < 0.01 | 0.99 | < 0.01 | 0.50 | **-0.14** | **< 0.01\*\*** |
| Breathing anxiety | 1.44 | 0.02 | 0.16 | 0.85 | -1.77 | 0.77 |
| Breathing difficulty | 0.59 | 0.05 | -0.74 | 0.08 | 5.49 | 0.08 |

**Supplementary Table 7:** Related to Figure 6. Correlation matrix across task modalities, with Pearson’s (A) and Spearman’s (B) correlation coefficients given above the diagonal and p values below the diagonal. Correlations with a p value<0.05 are represented in bold text, and those p<0.01 are shaded grey. Variables: 1) State anxiety (STAI-S); 2) Anxiety disorder (GAD-7); 3) Anxiety sensitivity (ASI); 4) Depression (CES-D); 5) Body perception (BPQ); 6) Interoceptive awareness (MAIA); 7) Breathing-related catastrophising (PCS-B); 8) Breathing-related vigilance (PVQ-B); 9) Perceptual threshold (from the FDT); 10) Decision bias (from the FDT); 11) Metacognitive bias (average confidence, from the FDT); 12) Metacognitive performance (from the FDT); 13) BOLD activity associated with positive predictions (from the BLT); 14) BOLD activity associated with negative predictions (from the BLT); 15) BOLD activity associated with positive prediction errors (from the BLT); 63) BOLD activity associated with negative prediction errors (from the BLT).

| A) CORRELATION MATRIX: PEARSON |  |  |  |  |  |  |  |  |  |  |  |  |  |  |  |  |
| --- | --- | --- | --- | --- | --- | --- | --- | --- | --- | --- | --- | --- | --- | --- | --- | --- |
|  | 1 | 2 | 3 | 4 | 5 | 6 | 7 | 8 | 9 | 10 | 11 | 12 | 13 | 14 | 15 | 16 |
| 1 |  | **0.70** | **0.57** | **0.72** | 0.22 | **-0.57** | **0.42** | 0.06 | **0.26** | **-0.28** | **-0.30** | -0.23 | -0.04 | 0.10 | 0.01 | -0.10 |
| 2 | **<0.01** |  | **0.55** | **0.72** | **0.35** | **-0.51** | **0.39** | 0.12 | 0.18 | -0.16 | -0.25 | -0.17 | -0.07 | 0.04 | 0.12 | -0.15 |
| 3 | **<0.01** | **<0.01** |  | **0.68** | **0.40** | **-0.40** | **0.62** | 0.20 | 0.20 | -0.18 | **-0.30** | -0.21 | 0.01 | 0.13 | 0.16 | -0.03 |
| 4 | **<0.01** | **<0.01** | **<0.01** |  | **0.31** | **-0.43** | **0.61** | **0.27** | **0.28** | -0.14 | -0.24 | -0.17 | -0.01 | 0.13 | 0.12 | -0.10 |
| 5 | 0.10 | **0.01** | **<0.01** | **0.02** |  | -0.16 | 0.25 | -0.02 | 0.04 | 0.19 | -0.03 | -0.12 | 0.06 | 0.06 | 0.06 | -0.08 |
| 6 | **<0.01** | **<0.01** | **<0.01** | **<0.01** | 0.24 |  | **-0.34** | 0.02 | -0.13 | 0.07 | 0.23 | 0.16 | 0.14 | -0.01 | -0.12 | 0.07 |
| 7 | **<0.01** | **<0.01** | **<0.01** | **<0.01** | 0.05 | **0.01** |  | **0.49** | -0.01 | 0.01 | -0.21 | **-0.30** | 0.17 | 0.24 | -0.05 | -0.10 |
| 8 | 0.66 | 0.36 | 0.13 | **0.04** | 0.89 | 0.87 | **<0.01** |  | -0.10 | -0.05 | 0.10 | 0.06 | 0.16 | 0.13 | 0.09 | -0.08 |
| 9 | **0.05** | 0.17 | 0.13 | **0.03** | 0.78 | 0.34 | 0.97 | 0.48 |  | 0.20 | 0.05 | 0.02 | -0.12 | -0.03 | 0.03 | -0.10 |
| 10 | **0.04** | 0.23 | 0.18 | 0.29 | 0.16 | 0.62 | 0.95 | 0.69 | 0.13 |  | **0.35** | 0.02 | -0.06 | 0.01 | 0.14 | 0.06 |
| 11 | **0.02** | 0.06 | **0.02** | 0.07 | 0.85 | 0.08 | 0.12 | 0.46 | 0.70 | **0.01** |  | **0.35** | -0.21 | **-0.28** | -0.00 | 0.07 |
| 12 | 0.09 | 0.20 | 0.11 | 0.20 | 0.38 | 0.22 | **0.02** | 0.65 | 0.87 | 0.87 | **0.01** |  | -0.23 | -0.23 | 0.13 | **0.42** |
| 13 | 0.75 | 0.60 | 0.91 | 0.96 | 0.64 | 0.29 | 0.19 | 0.22 | 0.36 | 0.68 | 0.12 | 0.08 |  | **0.79** | -0.22 | **-0.30** |
| 14 | 0.44 | 0.76 | 0.31 | 0.34 | 0.68 | 0.97 | 0.07 | 0.32 | 0.83 | 0.94 | **0.03** | 0.08 | **<0.01** |  | -0.10 | **-0.46** |
| 15 | 0.92 | 0.39 | 0.23 | 0.37 | 0.64 | 0.36 | 0.72 | 0.52 | 0.80 | 0.28 | 0.98 | 0.33 | 0.10 | 0.45 |  | 0.08 |
| 16 | 0.45 | 0.26 | 0.84 | 0.46 | 0.53 | 0.58 | 0.46 | 0.57 | 0.45 | 0.63 | 0.60 | **<0.01** | **0.02** | **<0.01** | 0.57 |  |
| B) CORRELATION MATRIX: SPEARMAN |  |  |  |  |  |  |  |  |  |  |  |  |  |  |  |  |
|  | 1 | 2 | 3 | 4 | 5 | 6 | 7 | 8 | 9 | 10 | 11 | 12 | 13 | 14 | 15 | 16 |
| 1 |  | **0.71** | **0.55** | **0.77** | **0.26** | **-0.57** | **0.34** | 0.05 | 0.24 | -0.18 | -0.22 | -0.23 | -0.18 | 0.06 | 0.07 | -0.03 |
| 2 | **<0.01** |  | **0.53** | **0.67** | **0.40** | **-0.51** | **0.27** | 0.10 | 0.13 | -0.18 | **-0.27** | -0.13 | -0.12 | 0.01 | 0.15 | -0.05 |
| 3 | **<0.01** | **<0.01** |  | **0.67** | **0.47** | **-0.36** | **0.58** | 0.22 | 0.19 | -0.15 | **-0.29** | -0.14 | -0.10 | 0.13 | 0.22 | 0.04 |
| 4 | **<0.01** | **<0.01** | **<0.01** |  | **0.37** | **-0.40** | **0.51** | 0.19 | 0.24 | -0.22 | **-0.26** | -0.13 | -0.07 | 0.11 | 0.21 | 0.01 |
| 5 | **0.05** | **<0.01** | **<0.01** | **<0.01** |  | -0.17 | **0.26** | 0.03 | 0.05 | 0.08 | -0.05 | -0.01 | -0.01 | 0.01 | 0.20 | 0.01 |
| 6 | **<0.01** | **<0.01** | **0.01** | **<0.01** | 0.20 |  | **-0.28** | -0.00 | -0.11 | 0.04 | 0.15 | 0.08 | 0.03 | -0.07 | -0.15 | 0.03 |
| 7 | **0.01** | **0.04** | **<0.01** | **<0.01** | **0.05** | **0.04** |  | **0.56** | -0.01 | 0.03 | -0.24 | **-0.29** | 0.17 | 0.22 | -0.06 | 0.02 |
| 8 | 0.71 | 0.47 | 0.10 | 0.16 | 0.82 | 0.98 | **<0.01** |  | -0.08 | -0.11 | 0.03 | -0.04 | 0.17 | 0.09 | 0.10 | -0.06 |
| 9 | 0.07 | 0.35 | 0.16 | 0.07 | 0.73 | 0.42 | 0.96 | 0.54 |  | 0.14 | 0.13 | 0.02 | -0.10 | 0.06 | -0.03 | -0.14 |
| 10 | 0.18 | 0.17 | 0.26 | 0.10 | 0.57 | 0.79 | 0.82 | 0.40 | 0.30 |  | **0.34** | 0.07 | -0.08 | -0.10 | 0.07 | 0.12 |
| 11 | 0.10 | **0.04** | **0.03** | **0.04** | 0.68 | 0.27 | 0.07 | 0.81 | 0.34 | **0.01** |  | 0.32 | -0.10 | -0.18 | -0.00 | 0.07 |
| 12 | 0.08 | 0.32 | 0.31 | 0.31 | 0.92 | 0.55 | **0.03** | 0.77 | 0.90 | 0.59 | **0.01** |  | -0.20 | -0.23 | **0.30** | **0.47** |
| 13 | 0.18 | 0.38 | 0.45 | 0.63 | 0.92 | 0.84 | 0.19 | 0.20 | 0.44 | 0.55 | 0.47 | 0.14 |  | **0.62** | -0.21 | -0.08 |
| 14 | 0.67 | 0.92 | 0.33 | 0.42 | 0.97 | 0.60 | 0.10 | 0.51 | 0.67 | 0.47 | 0.18 | 0.08 | **<0.01** |  | -0.19 | **-0.32** |
| 15 | 0.59 | 0.25 | 0.09 | 0.12 | 0.13 | 0.27 | 0.66 | 0.44 | 0.81 | 0.63 | 0.98 | **0.02** | 0.11 | 0.16 |  | 0.20 |
| 16 | 0.82 | 0.69 | 0.78 | 0.95 | 0.97 | 0.84 | 0.89 | 0.66 | 0.31 | 0.37 | 0.62 | **<0.01** | 0.54 | **0.01** | 0.12 |  |

